# Sub-nucleosomal organization in urine cell-free DNA

**DOI:** 10.1101/696633

**Authors:** Havell Markus, Jun Zhao, Tania Contente-Cuomo, Elizabeth Raupach, Ahuva Odenheimer-Bergman, Sydney Connor, Bradon R. McDonald, Elizabeth Hutchins, Marissa McGilvery, Michelina C. de la Maza, Kendall Van Keuren-Jensen, Patrick Pirrotte, Ajay Goel, Carlos Becerra, Daniel D. Von Hoff, Scott A. Celinski, Pooja Hingorani, Muhammed Murtaza

## Abstract

Cell-free DNA (cfDNA) in urine is a promising analyte for noninvasive diagnostics. However, urine cfDNA is highly fragmented and whether characteristics of these fragments reflect underlying genomic architecture is unknown. Here, we perform comprehensive characterization of fragmentation patterns in urine cfDNA. We show modal size and genome-wide distribution of urine cfDNA fragments are consistent with transient protection from degradation by stable intermediates of nucleosome disassembly. Genome-wide nucleosome occupancy and fragment sizes in urine cfDNA are informative of cell of origin and renal epithelial cells are amongst the highest contributors in urine. Compared to a nucleosome occupancy map based on control urine samples, we observe a higher fraction of fragments with aberrant ends in cancer patients, distinguishing cancer samples with an area under the curve of 0.89. Our results demonstrate sub-nucleosomal organization in urine cfDNA and are proof-of-principle that genome-wide fragmentation analysis of urine cfDNA can enable cancer diagnostics.

---

Circulating cell-free DNA (cfDNA) has emerged as an informative biomarker in prenatal, organ transplant and cancer patients. Recent studies have shown that genome-wide distribution and fragmentation of cfDNA in plasma is not random. Plasma cfDNA fragments have a modal size of 167 bp, are protected from degradation within mono-nucleosomes and their positioning captures nucleosome footprints of contributing tissues^1^. In cancer patients, these observations potentially enable cancer detection^2^, inference of tissue of origin^3^ and inference of gene expression^4^. In addition, deviations from expected fragment size and positioning can be leveraged to improve signal-to-noise ratio for somatic genomic alterations in plasma cfDNA^5^.

Collection of blood plasma requires venipuncture and plasma volume obtainable at a single time point is limited. In contrast, urine can be collected noninvasively, with minimal assistance and in larger volumes. However, there has been limited success in diagnostic development using urine cfDNA so far. There are multiple reports that cfDNA fragments are more degraded, shorter and variably sized in urine compared to plasma^6, 7^, impeding targeted analysis of genomic alterations. Comprehensive characterization of fragment sizes and positioning in urine cfDNA has not been reported and whether any genome-wide organization is preserved is unknown.

We characterized fragmentation patterns in urine cfDNA using whole genome sequencing. To our surprise, urine samples from healthy volunteers predominantly showed a modal size of 80-81 bp, suggesting non-random cfDNA fragmentation in urine. Here, we evaluate this hypothesis and investigate fragment size, distribution and nucleosome positioning in urine cfDNA. We describe correlation between cfDNA fragmentation patterns in urine and chromatin accessibility as well as gene expression in contributing cells. In cancer patients, we report a framework to leverage genome-wide differences in urine cfDNA fragmentation as a diagnostic approach.

## Results

To investigate fragment size distribution with high resolution and compare with plasma samples, we performed whole genome sequencing (WGS) of 30 urine and 15 plasma cfDNA samples collected from unrelated healthy volunteers (mean physical coverage of 7x and total coverage 196x in urine, mean 3.6x and total coverage 58x in plasma). In plasma cfDNA, we observed a modal fragment size of 167 bp, as reported previously^8^ (Fig. 1a, Supplementary Fig. 1). In urine cfDNA, we found the modal fragment size in 23/30 samples was 80-81 bp. In an additional 6/30 samples, the modal fragment size was 111-112 bp (Fig. 1b, Supplementary Fig. 2). In both urine and plasma, we found a 10 bp step pattern but the amplitude of each fragment size peak was much greater in urine (Fig. 1c). While the size distribution of plasma cfDNA showed one predominant peak, fragment size peaks in urine cfDNA were more evenly distributed relative to the mode.

**Figure 1:**
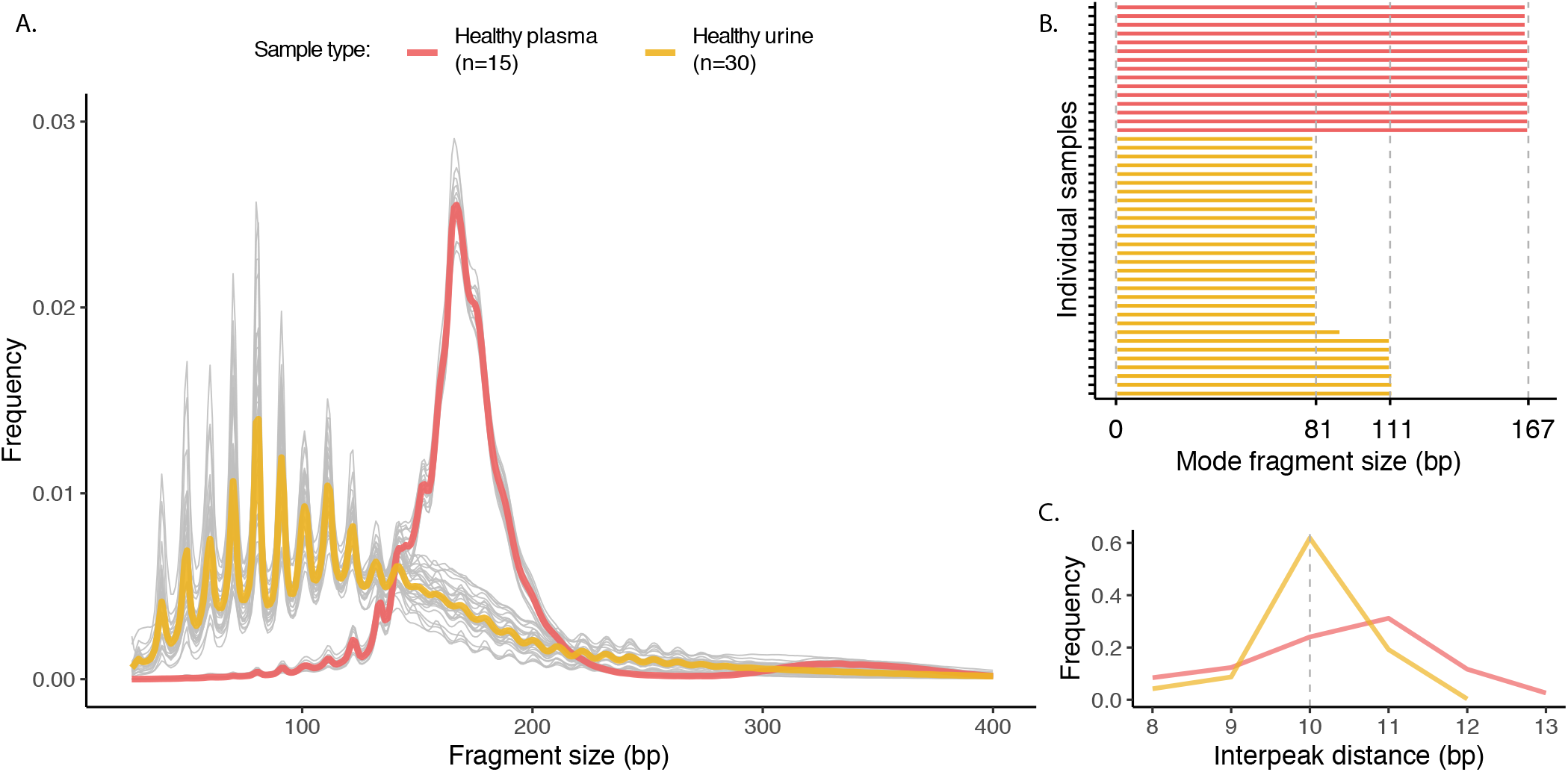
Comparison of fragment size between plasma and urine samples. (A) Genome-wide fragment size distributions were measured. Individual control plasma and urine samples are shown as grey lines. Mean across plasma and urine samples are shown as red and yellow lines, respectively. (B) Modal size in individual control plasma and urine samples was defined as the fragment size with the highest frequency. (C) Interpeak (peak-to-peak) distance of periodic peaks for plasma and urine samples.

We hypothesized that a modal size of 80-81 bp in the majority of samples may be associated with a stable intermediate product of histone-DNA interaction and nucleosome assembly. In vitro studies show that a histone H3-H4 tetramer is the most energetically favorable intermediate component during stepwise nucleosome assembly. The intermediate tetrasome binds a ~71 bp central region of the DNA originally wrapped in each mono-nucleosome^9^. To evaluate whether a similar mechanism may explain fragment sizes in urine cfDNA, we compared physical sequencing coverage between plasma and urine in a genomic region previously reported to have consistent strongly positioned nucleosomes^10^. In plasma and urine cfDNA, we found periodic peaks in coverage that were consistent in positioning between the two sample types (Fig. 2a, Supplementary Fig. 3). Individual urine peaks were narrower and occupied the center of corresponding plasma peaks (Fig. 2b). We investigated whether fragmentation sites in urine cfDNA were similar to those found in plasma. Using a plasma-based nucleosome occupancy map published previously^3^, we evaluated distance between fragment start or end sites, and centers of corresponding nucleosome dyad peaks in plasma and urine samples across 12.9 million nucleosomes. As expected, distribution of fragment start and end site distances in plasma showed distinct modes at 78 bp upstream and downstream of nucleosomal dyad peaks and reduced representation within the nucleosome core (Fig. 2c). In contrast, fragment start and end site distances in urine showed no distinct modes but a decreasing abundance of fragmentation sites as distance to the nucleosome dyad decreased. These results suggest nucleosomes dissociate under physiological conditions in urine and cfDNA becomes more vulnerable for enzymatic degradation but may be transiently protected by nucleosome assembly intermediates, predominantly by association with H3-H4 tetramers.

**Figure 2:**
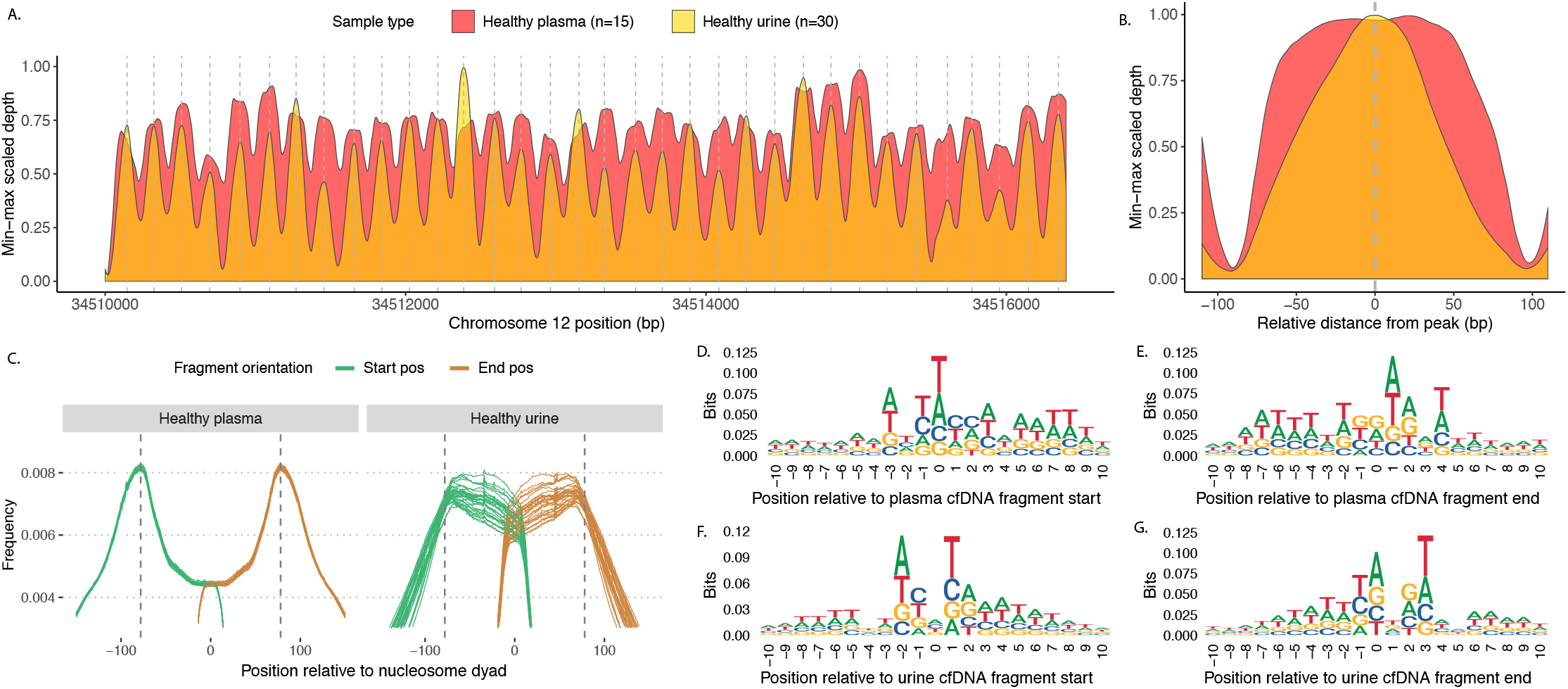
Comparison of sequencing coverage and fragment end sites between plasma and urine samples. (A) LOESS smoothed and min-max scaled physical sequencing coverage of pooled plasma and urine samples in a genomic region with stable nucleosomes (Chromosome 12p11.1). The vertical grey lines depict the local maxima of each peak for the pooled urine samples. (B) Mean smoothed physical sequencing coverage calculated by centering all peaks at the local maxima. (C) Genome-wide distribution of fragment start and end sites of individual plasma and urine samples relative to nucleosome dyads inferred from a published plasma-based nucleosome occupancy map. The vertical lines are drawn at 78 bp downstream and upstream from the nucleosome dyad represent the theoretical boundary of nucleosome octamer. (D-F) Nucleotide frequencies surrounding 10 bp upstream and downstream of fragment start and end sites in pooled plasma and urine.

To investigate whether histones are detectable in urine, we evaluated commerically available pooled control urine samples using proteomics analysis. Proteins were extracted using trichloroacetic acid percipitation to enrich for highly basic proteins. Peptides were generated by trypsin digestion for mass spectrometry analysis using LC-MS/MS and offline fractionation. This analysis yielded 4806 unique peptides (False Discovery Rate, FDR<0.01) and 1374 unique proteins (FDR<0.05). We detected 2-5 high confidence unique peptides from all four nucleosomal histone proteins (H2A, H2B, H3, and H4) and histone H1 with percent protein coverage ranging from 9%-42% (Supplementary Table 1). High confidence peptides were detected with 2-19 peptide spectral matches. The presence of histones in urine is consistent with a model that urine cfDNA may be transiently protected from degradation by association with histone proteins. Detection of all canonical histone proteins and a range of DNA fragment sizes suggests cfDNA in urine may be heterogeneous in its packaging including a mixture of tetrasome and hexasome structures.

We also compared nucleotide frequencies observed in cfDNA fragment ends between urine and plasma. As reported recently^11^, we found a palindromic pattern of per base nucleotide frequencies 10 bp upstream and downstream of fragment start and end sites in plasma cfDNA, conserved across all 15 plasma samples (Fig. 2d, e, Supplementary Fig. 4, 5). In urine cfDNA, we found a different pattern from plasma which was also conserved across all 30 urine cfDNA samples (Fig. 2f, g, Supplementary Fig. 6, 7). To evaluate whether these sequence preferences vary for different fragment lengths, we divided fragments into bins by fragment size (55-65 bp, 66-75 bp, 76-85 bp and so on)^12^. In both plasma and urine, these patterns were conserved regardless of fragment size (Supplementary Fig. 8). These observations indicate that there may be different enzymes responsible for DNA degradation in plasma and urine. Alternatively, the same enzyme may have different sequence preferences under different physiological conditions in urine and plasma. A recent study suggests that DNase1-like3 is the predominant enzyme degrading chromatin in plasma^13^. In contrast, DNase1, an enzyme highly active in urine^14^, has preferential activity for naked DNA after DNA-bound proteins are removed^15^.

We next investigated whether urine cfDNA provided any insights into genome-wide nucleosome positioning. We pooled data from plasma and urine samples to generate nucleosome maps and identified 11.8 and 7.2 million nucleosome peaks respectively. We found 69.4% and 60.4% nucleosome peaks in plasma and urine maps overlapped with the most well characterized plasma-based nucleosome occupancy map published so far (Fig. 3a, b)^3^. Non-overlapping peaks had significantly lower confidence scores compared to overlapping peaks (Supplementary Fig. 9). The modal distance between consecutive adjacent nucleosome peaks in plasma and urine samples was 184 bp and 177 bp respectively, similar to earlier results and consistent with periodic nucleosomal positioning (Fig. 3c). When any two plasma-based nucleosome maps were compared, the distance between corresponding peaks was predominantly zero. In contrast, a comparison between urine and plasma nucleosome maps showed a wider spread of distances, suggesting differential positioning of nucleosomes in urine (Fig. 3d). Relative to the nucleosome occupancy map based on urine instead of plasma, we found distinct modes in the distribution of start and end sites for urine cfDNA fragments around the nucleosome center (Fig. 2c, Supplementary Fig. 10).

**Figure 3:**
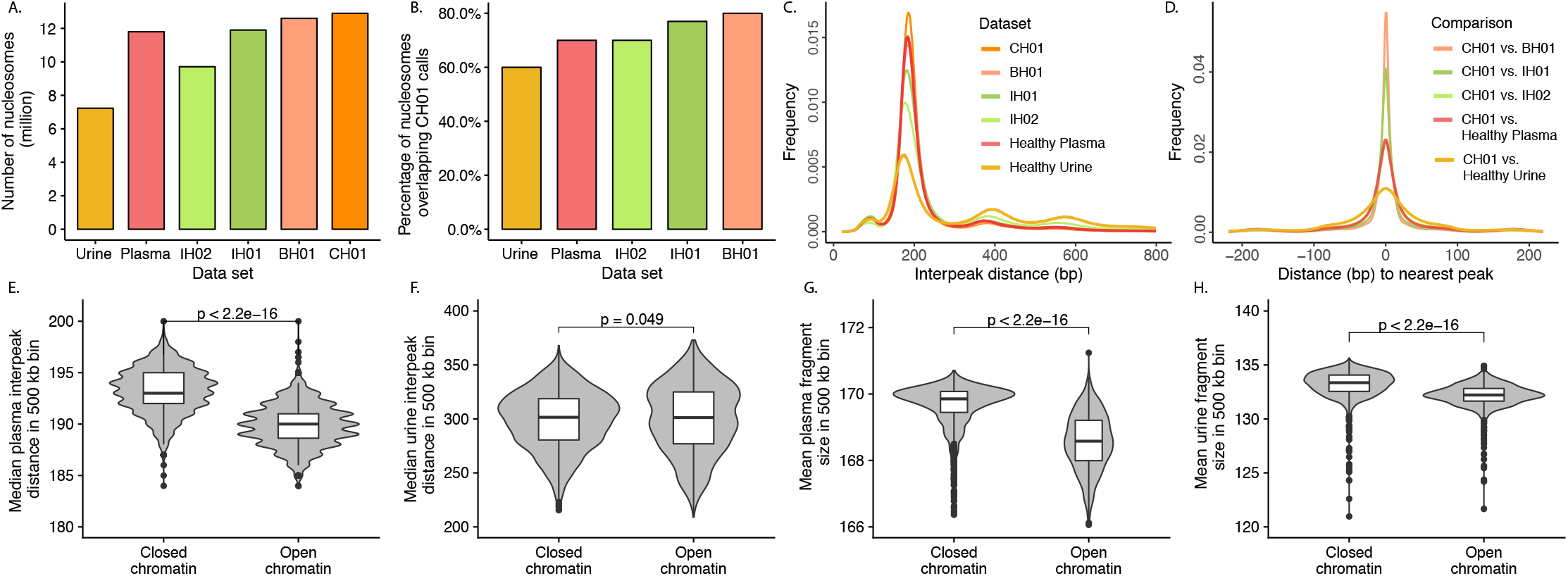
Benchmarking of a nucleosome occupancy map based on urine cfDNA. (A) Number of nucleosome calls inferred using window protection score method for pooled urine and plasma samples, compared with published plasma-based nucleosome tracks (IH01, IH02, BH01, and CH01). (B) Percentage of nucleosome calls from pooled urine, pooled plasma, IH01, IH02, and BH01 nucleosome tracks overlapping the CH01 nucleosome track. (C) Distance between adjacent peak centers (interpeak distance) in each nucleosome track. (D) Pairwise comparison of peak calls between CH01 (plasma) and all other tracks. The distribution of distances between corresponding peak centers are shown. Negative and positive distances indicate the nearest peak centers in CH01 are downstream or upstream, respectively. (E-F) Comparison of plasma and urine median interpeak distance in 500 kb bins annotated as compartment B (closed chromatin regions) and compartment A (open chromatin regions) from Hi-C chromatin contact map of a lymphoblastoid cell line (GM12878). (G-H) Comparison of plasma and urine mean fragment size in 500 kb bins annotated as compartment A and B.

To further assess if urine cfDNA fragments are informative of genome-wide nucleosome positioning in cells, we compared interpeak distances within open and closed chromatin regions. We considered non-overlapping windows of 500 kb across the genome annotated using published Hi-C chromatin contact maps for a lymphoblastoid cell line (GM12878) as compartment A (transcriptionally active and enriched for open chromatin) and compartment B (transcriptionally silent and enriched for closed chromatin). In agreement with earlier results, plasma samples showed significantly smaller distances between adjacent nucleosomes on average in compartment A than compartment B (p<2×10^−16^, Student’s t-test; Fig. 3e)^16^. However, a similar trend was not observed in urine samples (Fig. 3f), likely due to a sparser set of nucleosome calls available from urine and reliance on a lymphoblastoid cell line for compartment A/B annotation. As an alternative approach, we investigated whether cfDNA fragments are more degraded and shorter in size in open chromatin regions. We observed median fragment sizes in compartment A (open chromatin regions) were significantly shorter than compartment B (closed chromatin regions) in both plasma and urine samples (p<2×10^−16^, Student’s t-test; Fig. 3g, h).

Given a non-random distribution of urine cfDNA fragments across the genome, we hypothesized that genome-wide fragmentation patterns and positioning in urine may be informative of tissue of origin. Based on differences observed in urine cfDNA fragment size in open and closed chromatin regions, we developed a computational approach for correlating fragment size differences between active and inactive genomic regions to DNase I hypersensitive sites (DHS) across 116 cell types and tissues (using a published dataset)^17–19^. For each individual sample, we calculated median fragment size within non-overlapping 500 kb windows for all autosomes and normalized all median values to z-scores. Windows with negative and positive z-scores (shorter and longer fragments) associated with compartments A and B (open and closed chromatin) of GM12878 respectively (Fig. 4a, b, c). This association was stronger for plasma samples than urine samples (cosine similarity of 0.53 and 0.37 respectively). For each cell line, we calculated the number of DHS regions annotated in non-overlapping 500 kb windows for all autosomes and normalized all the counts to z-scores. Using these two sets of z-scores, we calculated the cosine similarity between the fragment size vector from each individual sample and the DHS vector for each cell line. In open chromatin regions, we expect a fewer number of DHS sites and shorter cfDNA fragments (a positive correlation). For pooled plasma, we found the highest cosine similarity with lymphoid or myeloid cells (Fig. 4d, Supplementary Fig. 11a, Supplementary Data 1). In contrast, for pooled urine, we found the highest cosine similarity with epithelial, renal epithelial and renal cortical cells (Fig. 4e, Supplementary Fig. 11b). The mean quantile normalized cosine similarity (MQNCS) for lymphoid or myeloid cells (n=21) was higher in plasma samples compared to urine samples (p<0.001, Student’s t-test). Conversely, MQNCS for renal cells (n=4) was lower in plasma samples compared to urine samples (p<0.01, Student’s t-test; Fig. 4f, g). These results suggest renal and uro-epithelial cells contribute a large fraction of cfDNA in urine. In urine samples, cell-type specific MQNCS scores were much more variable compared to plasma samples, suggesting that tissue contributions in urine samples are more variable.

**Figure 4:**
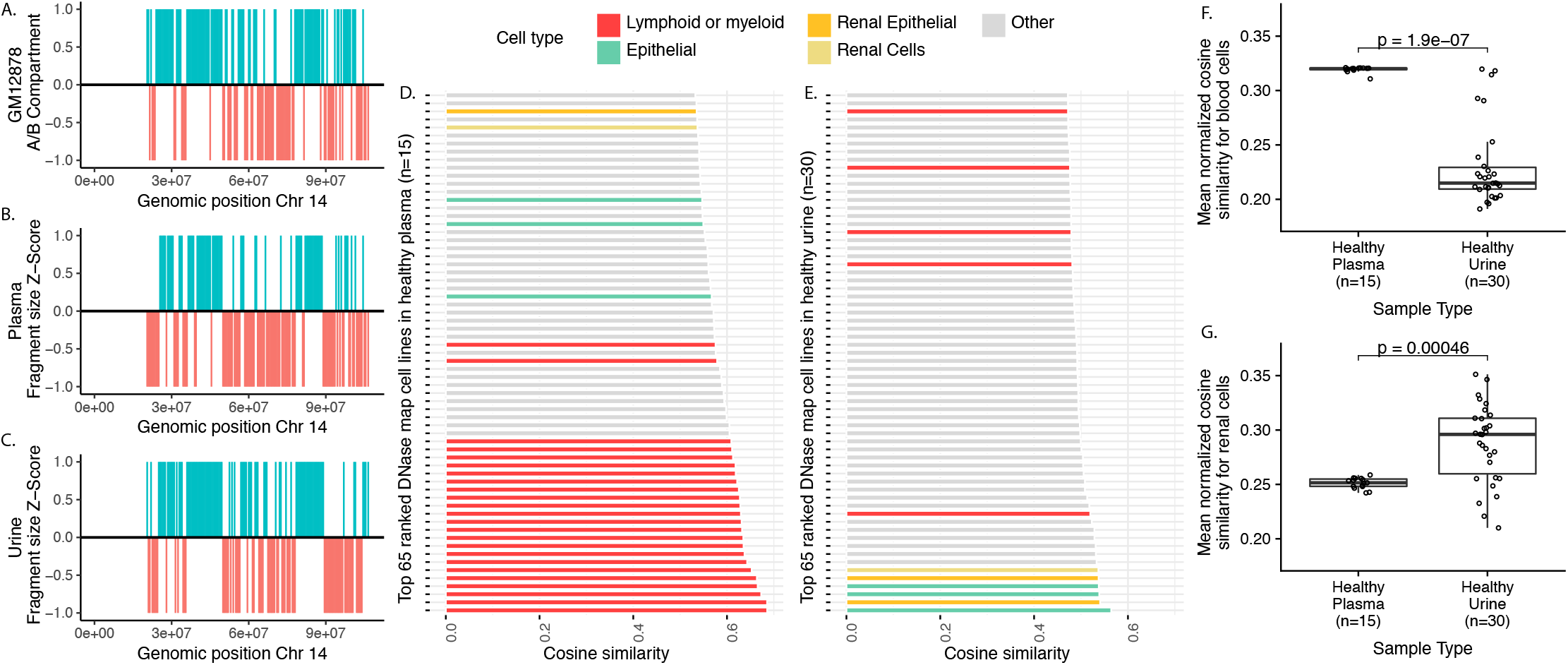
Comparison of cfDNA fragment size with chromatin accessibility across cell types. (A) Distribution of A (open chromatin) and B (closed chromatin) compartments in non-overlapping 500 kb bins of Chr 14 from Hi-C chromatin contact map of lymphoblastoid cell lines (GM12878). A and B compartments are shown in red and blue colors respectively. (B-C) Distribution of median cfDNA fragment size in non-overlapping 500 kb bins of Chr 14 normalized to a z-score for pooled plasma and urine samples respectively. Bins with negative and positive z-score values were converted to −1 and 1 and colored red and blue respectively. (D-E) 65 cell lines or tissues with highest cosine similarity between cfDNA fragment size and DHS sites in 500 kb bins across the genome. (F) Comparison of mean quantile normalized cosine similarity scores for bone marrow, lymphoid or myeloid cell lines (n=24) in individual plasma and urine samples. (G) Comparison of mean quantile normalized cosine similarity scores for renal tissues and renal epithelial cell lines (n=4) for individual plasma and urine samples.

To validate tissue contribution in urine using an alternative approach, we evaluated cfDNA coverage around transcription start sites in urine and plasma. As previously reported, coverage in pooled plasma samples showed a nadir at ~150 bp upstream of transcription start sites (TSS) with greater loss of coverage for highly expressed genes (Fig. 5a)^4^. We observed a similar pattern in pooled urine samples (Fig. 5b). However, in contrast to plasma DNA, overall urine cfDNA coverage in the 2 kb region around TSS is higher than surrounding loci and coverage is higher downstream of TSS than upstream, particularly for highly expressed genes. To infer tissue of origin, we compared sequencing coverage at TSS in cfDNA to gene expression in 64 human cell lines and 37 primary tissues (Human Protein Atlas). For each plasma or urine sample, we measured coverage at the nucleosome-depleted region (NDR), from –150 bp to +50 bp around TSS of protein coding genes on autosomes. We calculated Spearman’s rank correlation coefficient (Spearman’s rho) between mean NDR coverage and individual gene expression values in each cell line or tissue (Supplementary Data 2). We expect a stronger negative correlation for cell types contributing greater amounts of cfDNA. As expected, pooled plasma showed the most negative correlations with lymphoid and myeloid cell lines or with bone marrow tissue (Supplementary Fig. 12a). Pooled urine was most negatively correlated with epithelial, lymphoid, myeloid, and embryonal kidney cell lines (Supplementary Fig. 12b). We evaluated the change in cell line and tissue ranks between plasma and urine. Lymphoid and myeloid cell lines and bone marrow tissue had the largest decrease while renal epithelial, urinary bladder, and epithelial cell lines had largest increase in rank in urine compared to plasma (Fig. 5c, Supplementary Fig. 13, Supplementary Data 3). The mean quantile normalized Spearman’s rho (MQNSR) for lymphoid and myeloid cell lines or bone marrow tissue (n=16) was lower in plasma compared to urine samples (p<0.001, Student’s t-test; Fig. 5d). Conversely, MQNSR for renal epithelial cell line and urinary bladder cell line was lower in urine compared to plasma samples (both p<0.001, Student’s t-test; Fig. 5e, f).

**Figure 5:**
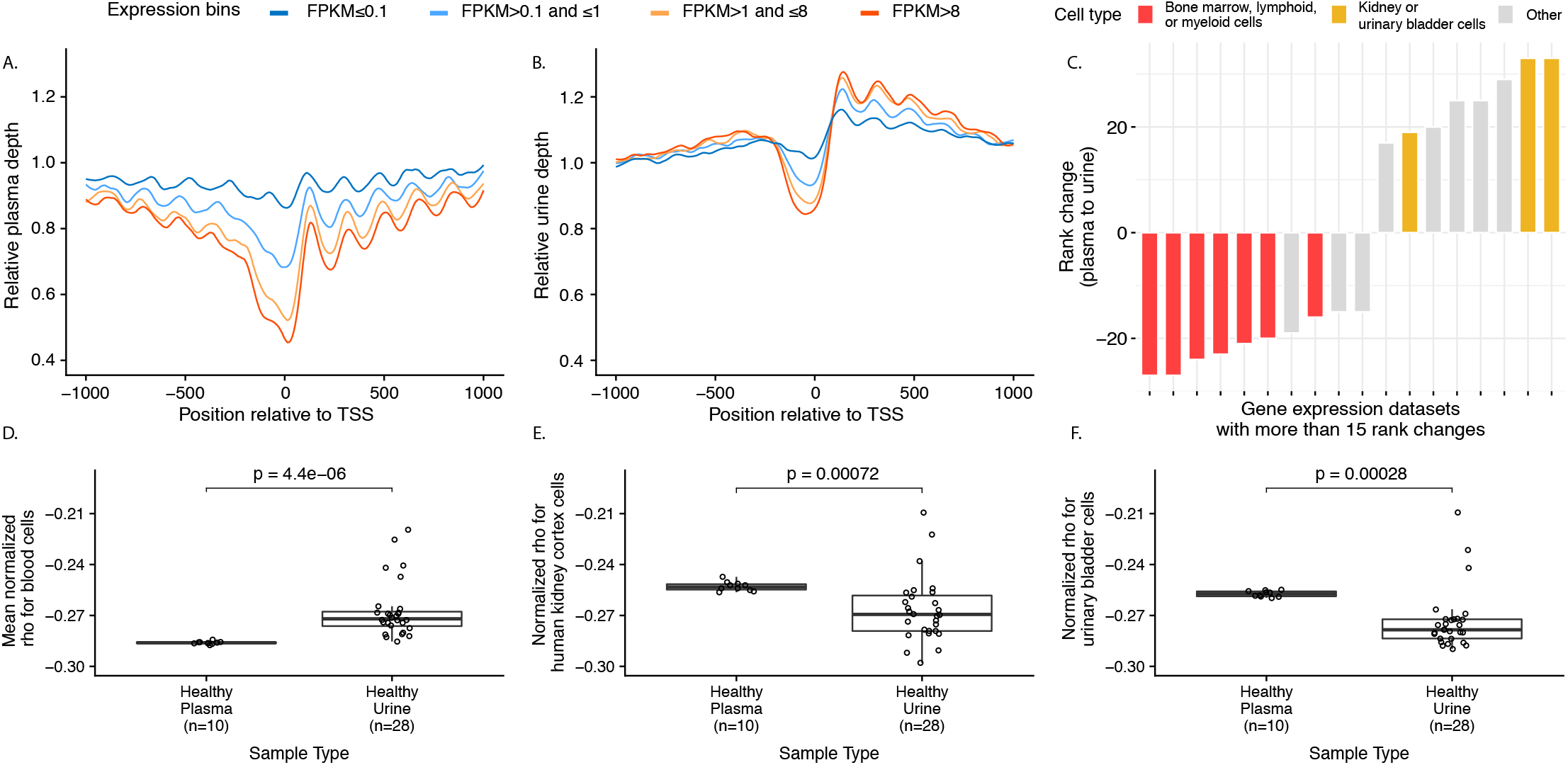
Comparison of cfDNA coverage at transcription start sites and correlation to gene expression across cell types. (A-B) Mean pooled plasma and urine sequencing depth at the transcription start sites (TSS) of genes binned by their expression levels in fragments per kilobase of transcript per million mapped reads (FPKM). A depletion of coverage is observed at transcription start sites in plasma (A) and in urine samples (B). Such depletion is greatest for genes with the highest expression. (C) Rank changes in correlation between sequencing coverage in the nucleosome depleted region and gene expression across plasma and urine cfDNA. Cell lines that changed by at least 15 ranks are shown here. (D-F) Comparison of mean quantile normalized Spearman’s rho for gene expression datasets from blood (n=16), renal cortex (n=1), and urinary bladder (n=1) in individual plasma and urine samples.

Our observations suggested a stable genome-wide distribution of urine cfDNA fragments that was informative of chromatin landscapes in the contributing cell types. To investigate if a deviation from expected cell types could be detected in cancer patients, we performed WGS on pre-treatment urine samples from 10 patients with pediatric solid cancers (mean physical coverage of 21.6x) and 12 patients with pancreatic cancer (mean physical coverage of 0.72x).

Urine cfDNA fragment size distribution was consistent with that observed in healthy volunteers (Supplementary Fig. 14). We built a reference nucleosome occupancy map using pooled data from 20/30 urine samples from healthy volunteers (12 females and 8 males, mean physical coverage of 155x). Against this map, we calculated the fraction of aberrant fragments starting or ending within 65 bp downstream or upstream of the nucleosome center in all urine samples including: 20/30 control samples used to create the map (training set of healthy volunteers), 10/30 additional control samples (test set of healthy volunteers), 10 pre-treatment samples from resectable and localized pediatric cancer patients, and 12 pre-treatment samples from patients with pancreatic adenocarcinoma (7 patients with stage I-II and 5 patients with stage IV disease; Fig. 6a, b, Supplementary Table 2, Supplementary Data 4). We observed no significant difference in fraction of aberrant fragments between the training and test sets, indicating that our reference map was comprehensive enough to capture variations in nucleosome positioning in urine cfDNA from healthy volunteers. In both sets of cancer patients, we found a significantly higher fraction of aberrant fragments when compared to the training samples (p<0.01, Student’s t-test), suggesting contribution of cfDNA from unexpected cell types with differences in genomic organization not captured by healthy volunteers. To explore this using an alternative approach, we investigated nucleotide frequencies around fragment starts and ends in individual samples to detect minor deviations between samples that may result from differences in chromatin accessibility across contributing cell types. We analyzed the per base mono-nucleotide frequencies in the 10 bp region upstream and downstream of fragment start and end sites (Supplementary Fig. 15, 16). Multidimensional scaling showed separation between healthy volunteers and pancreatic cancer patients in the third dimension (Fig. 6c). Using thresholds for fraction of aberrant ends and for multiple dimensions of nucleotide frequency at fragment ends, we evaluated the ability to distinguish urine samples from cancer patients from healthy volunteers. Using either feature individually or a combination of the two, we were able to distinguish cancer patients from healthy volunteers, achieving an area under the receiver operating characteristic curve of 85.3%-88.8% (Fig. 6d, Supplementary Fig 17). To evaluate whether aberrant fragments were tumor-specific, we compared fraction of urine cfDNA aberrant fragments between genomic regions known to be neutral, gained or lost due to somatic copy number changes in the tumor. In 4/6 patients, fraction of aberrant fragments was higher for genomic regions with copy number gains in tumor, compared to neutral and/or lost regions (one-tailed p<0.05, Student’s t-test; Supplementary Fig. 18, 20, 21, 22). In a fifth patient, no significant difference was observed (one-tailed p=0.079, Student’s t-test; Supplementary Fig. 19). In one patient, a reverse trend was observed but the tumor genome of this patient had widespread copy number changes with no clear baseline copy number neutral region (Supplementary Fig. 23).

**Figure 6:**
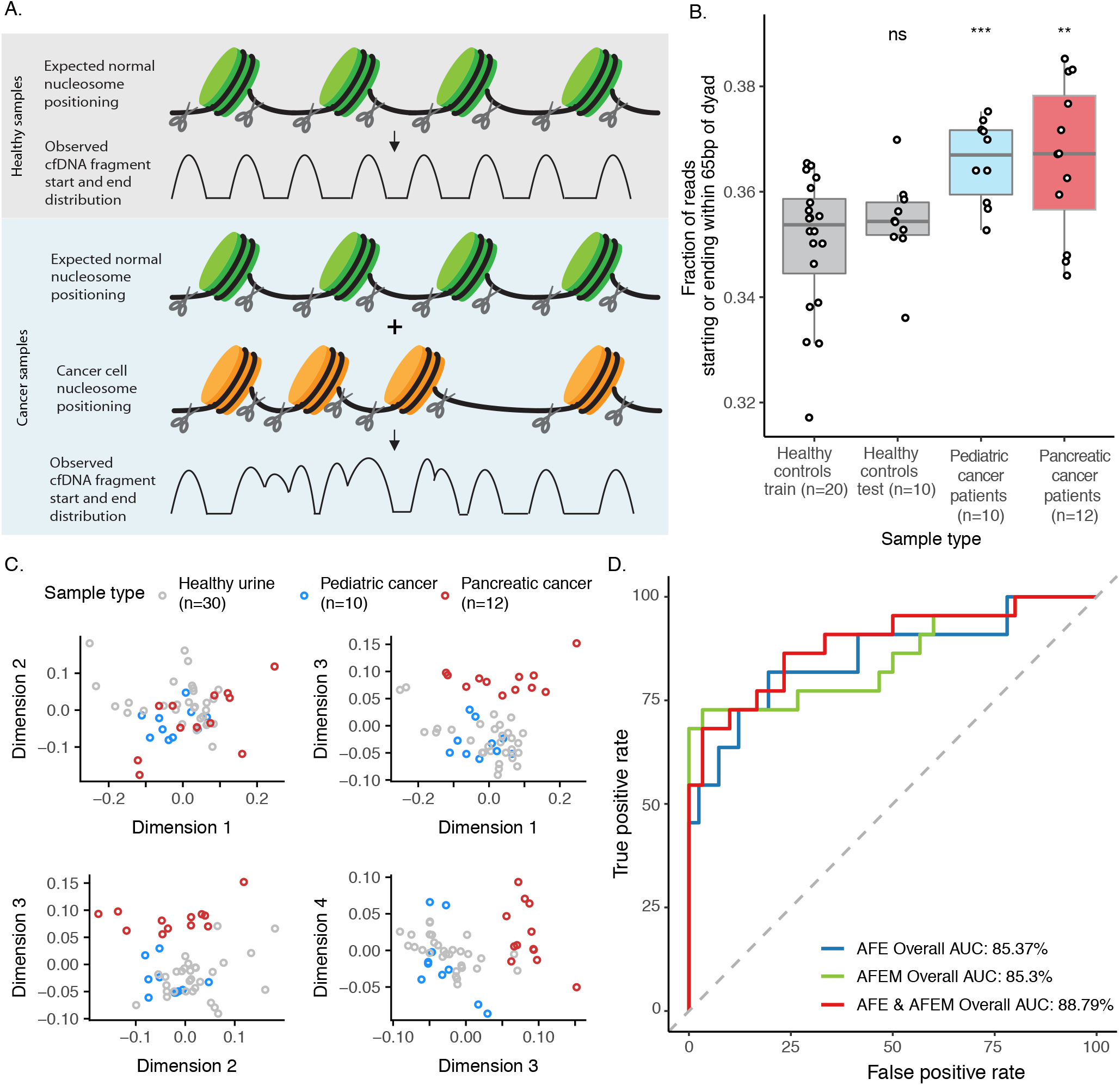
Evaluation of aberrant cfDNA fragments in urine from cancer patients. (A) Schematic representation of aberrant cfDNA fragments in urine samples from cancer patients. The illustration shows DNA wrapped in nucleosomes. Fragment start and end positions flank regions protected by nucleosomes and are clustered away from nucleosome centers. In patients with cancer, differences in nucleosome positioning in cancer cells that contribute cfDNA into urine may lead to a higher abundance of fragment start and end sites in unexpected genomic regions (such as regions protected by nucleosome in healthy control samples). (B) Fraction of urine cfDNA reads starting or ending within 65 bp of nucleosome dyads in reference nucleosome occupancy based on pooled urine cfDNA data from 20 controls (training set). The fractions from training set are compared to urine samples from 10 additional controls (test set), 10 pediatric cancer patients, and 12 pancreatic cancer patients. The ns, **, and *** represent p-values >0.05, <0.01, and <0.0001, respectively. (C) Multidimensional scaling (MDS) analysis of nucleotide frequencies in 20 bp region surrounding urine cfDNA fragment start and end sites (D) ROC analysis for classifying urine samples from controls and cancer patients using aberrant fragment ends (AFE), aberrant fragment end motifs (AFEM) or both. For AFE analysis, the fractions shown in (A) were used for ROC analysis. For AFEM and for the combination of AFE and AFEM, probabilities from a logistic regression fit to the first 4 MDS dimensions and AFE was used for ROC analysis.

## Discussion

Urine is a promising analyte for cancer-derived cell-free DNA to aid diagnostics and monitoring but this approach has been variably successful so far because urine cfDNA is highly degraded^20^. We found that fragment size distributions of urine cfDNA show recurrent modes at 81 and 111 bp and genome-wide distribution of urine cfDNA fragments is non-random. These observations mimic results from in vitro studies of nucleosome assembly^9^ and suggest transient protection of cfDNA fragments by association with subcomponents of unfolding nucleosomes in urine. We leveraged these features to produce a urine-based map of genome-wide nucleosome occupancy from healthy individuals and benchmarked it against similar plasma-based maps. We used associations between fragment size patterns and chromatin organization as well as transcription start site coverage and aggregate gene expression to identify cells and tissues contributing cfDNA into urine. Taken together, these results show a stable and reproducible genome-wide distribution of urine cfDNA fragments. They provide a framework for further development of urine-based liquid biopsies without reliance on genetic differences between somatic and germline, fetal and maternal, and donor and recipient genomes. In addition, the ability to infer and identify unexpected cell types contributing urine cfDNA may enable monitoring for complications or progression in other systemic diseases such as diabetes or hypertension.

In two different cohorts of cancer patients, we detected elevated fractions of aberrant fragments at unexpected genomic loci relative to a reference nucleosome occupancy map, suggesting contribution of urine cfDNA from unexpected cell types. This observation was further corroborated by deviations in nucleotide frequencies near cfDNA fragment ends in cancer patients compared to healthy controls. For a subset of patients where somatic copy number aberrations in the tumor were known, we found fractions of aberrant fragments were higher in genomic loci with copy number gains compared to neutral loci or those with copy number loss, suggesting our observations in urine cfDNA were driven by tumor cells. 17/22 cancer patients included in this study had non-metastatic and resectable disease and 19/22 patients had systemic solid tumors outside the genitourinary tract. Using fraction of aberrant reads and nucleotide frequencies in fragment ends, we achieved an area under the curve of 0.89 to distinguish cancer patients from healthy controls. These findings suggest a potential role for genome-wide analysis of urine cfDNA fragmentation and positioning in improving cancer diagnostics, particularly for early detection of cancer. Our findings also provide an opportunity to improve yield and accuracy for targeted somatic mutation assays by accounting for cfDNA degradation sites in the tissues of interest. This may require prior genome-wide characterization of urine samples from patients with advanced cancers of similar subtype and high tumor contribution in urine cfDNA.

In contrast to plasma, urine offers a truly noninvasive and abundant source of cancer-derived cell-free DNA. However, urine is not under as stringent homeostatic regulation and additional pre-analytical variation related to sample collection and processing must be considered. We observed a large variation in contribution of cfDNA from systemic and local tissues in urine samples, which may be explained by time elapsed since last void and duration that urine has incubated in the genitourinary system^6^. In addition, our approach for urine cfDNA is based on genome-wide fragmentation patterns and it lacks the inherent cancer specificity afforded by analysis of recurrent cancer-related somatic mutations. The cell types that contribute cfDNA can be affected by physiological states (differences in hydration status, pH or urea concentration), age, gender, co-morbidities such as diabetes and acute illnesses such as urinary tract infections. To utilize relative differences in fragmentation patterns for cancer detection, reference nucleosome occupancy maps based on urine samples from large cohorts of controls will be required. Both, pre-analytical and biological variation can also affect interpretation of tumor-derived fractions when using somatic mutation analysis in urine cfDNA from cancer patients^21^.

One limitation of this study is its reliance on urine samples from a limited number of controls and cancer patients (n=52). Although we have observed potential diagnostic value across two different cohorts of cancer patients, larger studies are needed to better refine diagnostic thresholds and estimate performance across cancer subtypes and disease stages. We have also relied on urine samples obtained from cancer patients at presentation and prior to treatment, when we expected tumor volume to be highest. Analysis of longitudinal samples obtained from cancer patients during treatment can provide more insight into the value of our approach for treatment monitoring. Another limitation is that we did not record or require specific time of day, hydration status, time since last void or other sources of biological and pre-analytical variation during urine collection. Our data suggest these factors should be accounted for in future studies.

In summary, our findings support a stable fragment distribution across the genome in urine cfDNA and set the stage for future investigation and development of urine-based diagnostic assays. We have shown proof-of-principle results that genome-wide fragmentation patterns and positioning in urine cfDNA yield diagnostic value for cancer patients. This approach can complement plasma-based liquid biopsy approaches for diagnosis and monitoring of cancer as well as non-malignant conditions.

## Methods

### Patients and samples

This study includes healthy volunteers enrolled at Translational Genomics Research Institute, Phoenix, AZ, USA under an approved IRB protocol number 20142638, pediatric cancer patients enrolled at Phoenix Children’s Hospital, Phoenix, AZ, USA under an approved IRB protocol number 16-141 and pancreatic cancer patients enrolled at Baylor Scott & White Research Institute under an approach IRB protocol number 015-196. Informed consent was obtained from all patients. For cancer patients, urine samples were collected at presentation and prior to treatment. Tumor samples analyzed were obtained at the time of diagnosis.

### Sample processing and cfDNA quantification in urine and plasma

Urine samples were processed within 1 hour of collection. We added 0.8 ml of 0.5 M EDTA to 40 ml of urine, centrifuged 10 ml aliquots at 1,600*g* for 10 min and stored at −80 °C. We extracted cfDNA from 10 ml urine using MagMAX Cell-Free DNA Isolation kit (Thermo Fisher Scientific) and eluted in 20-30 μl. Blood samples were collected in K2 EDTA BD Vacutainer tubes and processed within 2 hours of collection. Blood samples were centrifuged at 820*g* for 10 min at room temperature. 1 ml aliquots of plasma were further centrifuged at 16,000*g* for 10 min to pellet any remaining cellular debris. The supernatant was stored at −80 °C until DNA extraction. DNA was extracted using QIAamp Circulating Nucleic Acid kit (QIAGEN). We measured DNA yield using digital PCR assay^22^. In healthy volunteers, median urine cfDNA concentration was 0.82 ng/ml of urine (IQR: 2.3 ng/ml, n=30). Median plasma cfDNA concentration was 5.62 ng/ml of plasma (IQR: 4.75 ng/ml, n=16).

### Sequencing library preparation

For plasma cfDNA samples, we prepared whole genome sequencing libraries using 1 ng input from healthy volunteer samples using ThruPLEX Tag-seq (Takara Bio). We performed sequencing on HiSeq 4000 (Illumina) to generate 75 bp paired-end reads. The library prep kit introduces a 6 bp unique molecular identifier and an 8-11 bp random stem on both ends of DNA fragments. These tags were removed using a custom Python script. For urine cfDNA samples, we prepared whole genome sequencing libraries using 0.6-67.3 ng input using ThruPLEX Plasma-seq (Takara Bio). We performed sequencing on NovaSeq 6000 (Illumina) to generate 110 bp paired-end reads.

### Sequencing data and fragment size analysis

We de-multiplexed sequencing data based on sample specific barcodes and converted to fastq files using Picard tools v2.2.1 and using Illumina bcl2fastq v2.20.0.422 for plasma and urine data respectively, allowing 1 bp mismatch and requiring minimum base quality of 20. We aligned sequencing reads to the human genome using hg19 using bwa mem v0.7.15^23^. We sorted and indexed the bam files using samtools v1.3.1^24^. Reads with mapping quality <30 or unmapped, supplementary alignments, or not primary alignments were excluded from downstream analysis. Fragment size distribution and genomic coverage was calculated using Picard tools. One plasma sample was dropped from further analysis due to low coverage (<0.001x mean coverage). We calculated the modal fragment size and distance between fragment size peaks using a custom R script. We pooled plasma and urine controls by merging reads using samtools.

### Nucleosome coverage, fragment end position, and fragment end nucleotide frequencies

In a region with strongly positioned nucleosomes independent of tissue type^10^, we compared the physical coverage from pooled plasma and urine controls. For ease of visualization, we min-maxed (normalized data from 0 to 1) depth of coverage and applied a rough local polynomial regression fitting (LOESS) regression with a span of 0.02. Non-smoothed depth is shown in Supplementary Fig. 4. We calculated the mean smoothed physical coverage by centering all peaks in the region at their local maxima, estimated by inflection point.

To investigate the distance of fragment start and end sites in urine and plasma relative to their nearest nucleosome centers, we used a published plasma-based nucleosome occupancy map (CH01) as reference^3^. Paired end reads were summarized as fragments with their 3’ and 5’ position into a bed file using BEDTools v2.26.0^25^. Further analysis was carried out in R using the GenomicRanges package. The bed file of all fragments was intersected with the nucleosome occupancy track. For each overlap hit, the distance of fragment start and end position from the center of the respective nucleosome hit was calculated. In order to avoid fragments that might span more than one nucleosome, only 50-200 bp fragments were used for this analysis.

Using the fragments bed file, we created two additional bed files summarizing positions 10 bp upstream and downstream of fragment start and end site respectively. We extracted the genomic sequence of those regions using bedtools and calculated the mean per base mono-nucleotide frequencies using Homertools. We generated the start and end sequence motifs in R using the ggseqlogo package.

### Nucleosome map generation

We used published scripts based on window protection scores to create nucleosome occupancy maps using plasma and urine data^3^. For plasma samples, we used similar parameters as previously published: minimum fragment size of 120 bp, maximum fragment size of 180 bp, and window of 120 bp. For urine samples, we used the following parameters: minimum fragment size of 64 bp, maximum fragment size of 196 bp, and window of 120 bp. The parameters for urine cfDNA were changed to accommodate differences in fragment size distribution. To compare our plasma and urine maps with previously published tracks, we calculated the fraction of nucleosome calls that overlapped with CH01, the peak-to-peak distance between adjacent peaks (interpeak distance), and the distance to the nearest peak of CH01. The analysis was carried out in R using the GenomicRanges package.

### cfDNA characteristics in open and closed chromatin regions

We tiled all autosomes in the hg19 human genome into 500 kb non-overlapping bins. We excluded bins with mappability score <0.9, and bins within and or near the centromeric regions, resulting in 4,975 bins. We annotated each bin as compartment A (transcriptionally active and enriched for open chromatin) and compartment B (transcriptionally silent and enriched for closed chromatin) based on annotations from a previously published Hi-C chromatin contact map of lymphoblastoid cell lines (GM12878)^16^. We calculated median interpeak distance in each bin for the plasma and urine nucleosome maps using the GenomicRanges package. We calculated median fragment size in each bin using the Rsamtools package.

### Inference of tissue of origin by comparison with DNase Hypersensitive Sites

To infer tissue of origin for plasma and urine cfDNA, we calculated median fragment size (MFS) in each of the 500 kb bins and normalized the median values to a z-score (subtracting MFS in each bin to the mean MFS in all bins and dividing by the standard deviation of MFS in all bins). Bins with negative z-scores represent regions with higher fraction of shorter fragments, and bins with positive z-scores represent regions with higher fraction of longer fragments. We processed 116 DHS call sets of different cell lines published earlier in a similar manner^17^. The DHS data and annotations were downloaded from https://resources.altius.org/publications/Science_Maurano_Humbert_et_al/. For each call set, we calculated the number of DHS regions annotated in each 500 kb bin and normalized the counts to a z-score. Bins with negative z-scores represent regions with closed chromatin regions and bins with positive z-scores represent regions with open chromatin regions. We calculated the cosine similarity between the z-score vector for individual and pooled cfDNA samples and negative z-score vector for all DHS callsets. The cosine similarity between two vectors A and B can be calculated as: *A* · *B*/∥*A*∥∥ *B* ∥. To evaluate individual plasma and urine samples, we quantile normalized the cosine similarity (R preprocessCore package) in order to maintain both, cell line ranking and continuous nature of the metric. We calculated the mean quantile normalized cosine similarity (MQNCS) for all bone marrow, lymphoid or myeloid cell lines (n=24) and renal cell lines (n=4) for individual cfDNA samples.

### Inference of tissue of origin by comparison with gene expression

Using the previously generated cfDNA fragment bed files, we trimmed 61-800 bp fragments from both ends to contain 30 bp region downstream and upstream from the center (for odd fragment sizes we rounded the decimal down to closest integer). We left 20-60 bp fragments untrimmed. We converted the trimmed fragment bed files to bam files using bedtools. We calculated trimmed fragment coverage around the transcription start sites (TSS) of all genes in hg19 autosomes using the Rsamtools package. We normalized the coverage in TSS ± 1000 bp window around the TSS of all genes by mean depth in TSS – 3000 bp to TSS – 1001 bp and TSS + 1001 bp to TSS + 3000 bp regions. We further corrected the normalized coverage by the coding strand direction. We averaged the strand corrected normalized coverage around the TSS ± 1000 bp window across genes with similar gene expression values in plasma, as published earlier^4^. To infer tissue of origin using TSS coverage, we included samples with mean genomic coverage >3x (pooled plasma, pooled urine, 10 individual plasma controls, and 28 individual urine controls). We calculated the raw read depth coverage –150 bp to +50 bp around the TSS (Nucleosome Depleted Region coverage) for all genes in hg19 autosomes and correlated them to their respective expression values from 64 human cell lines and 37 primary tissues obtained from the Human Protein Atlas using Spearman’s rank correlation coefficient (Spearman’s rho). We also assessed the change in rank between pooled plasma and urine samples. To see whether this trend was consistent in individual plasma and urine samples, we quantile normalized the Spearman’s rho (R preprocessCore package) for all 64 human cell lines and 37 primary tissues across all individual samples (10 plasma and 28 urine) in order to maintain both rank and the continuous nature of the metric. We calculated the mean quantile normalized Spearman’s rho (MQNSR) for all bone marrow, lymphoid or myeloid tissues and cell lines (n=16) and two renal cell lines (RPTEC-TERT1 and RT4) for individual samples.

### Aberrant fragmentation ends and fragment end nucleotide frequency in cancer patients

We pooled reads from 20 healthy urine samples (12 females and 8 males) using samtools and built a urine reference nucleosome map (URNP) using parameters described earlier. We intersected individual fragment bed files with URNP using GenomicRanges package in R. For each overlap hit, the distance of fragment start and end position from the center of the respective nucleosome hit was calculated. We calculated the fraction of fragments that started or ended within 60 bp downstream or upstream of nucleosome center. These were counted as aberrant fragments, as they are being cleaved within or close to the nucleosome centers observed in reference samples. We compared the fraction of aberrant fragments in 20 control urine samples used to generate the URNP with urine samples from another 10 controls, 10 pediatric cancer patients, and 12 pancreatic cancer patients. We calculated the predictive performance of fraction of fragments with aberrant ends to distinguish between healthy and cancer samples using receiver operator curve (ROC) analysis (pROC R package). Since the fraction of fragments with aberrant ends in training and test control samples was similar, we used all 30 controls samples in the ROC analysis. We conducted ROC analyses on pediatric and pancreatic cancer samples separately and in combination.

Using the urine cfDNA fragment bed files from controls and cancer patients, we created two additional bed files summarizing positions 10 bp upstream and downstream of fragment start and end site respectively. We extracted the genomic sequence of those regions using bedtools and calculated the mean per base mono- and di-nucleotide frequencies as well as cumulative frequency of CpG, total G+C, total A+G, and total A+C using Homertools. For each individual sample, we summarized the various per base frequencies at fragment start and end sites in a single vector of length 168 in R. We concatenated the nucleotide frequency vector from all urine samples into one matrix (52 x 168). To reduce dimensions, we carried out Multidimensional Scaling (MDS) to reduce the data to 4 dimensions (52 x 4). We visualized whether nucleotide frequencies at fragment start and end sites could classify between healthy samples from cancer samples by plotting various combinations of the 4 dimensions. We refer to the nucleotide frequencies at fragment start and end sites as AFEM (aberrant fragment end motif). To calculate the predictive performance of AFEM to distinguish between healthy and cancer samples we fitted logistic regression using base *glm* function in R to the 4 MDS dimensions and used the predictive probability from the model to conduct ROC analysis. We conducted ROC analyses on pediatric and pancreatic cancer samples separately and in combination. We also combined the 4 MDS dimensions and AFE, and conducted an integrated ROC analysis using similar steps.

### Aberrant fragmentation ends in copy number aberration regions

To investigate whether the AFE was affected by underlying copy number changes in the tumor, we used data generated using exome sequencing of tumor and germline DNA samples from 2 patients with pediatric cancers and 4 patients with pancreatic cancer. We calculated regions with copy number aberrations using the R package Sequenza^26^. For each of the 6 patients, we marked the 4975 bins as copy number neutral, loss, or gain. We removed any bins that were partially segmented into two different copy number states. We calculated the AFE in each of the filtered 500 kb bins for urine samples from the 6 cancer patients and the 10 controls not used to build the URNP. For each patient, we calculated the AFE ratio in bin *i* as the AFE in bin *i* of patient sample divided by the mean AFE in bin *i* of the 10 healthy urine samples, as shown in Equation 1.

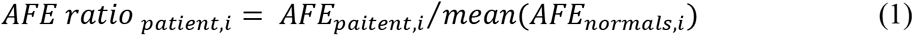

We also calculated background distribution of AFE ratio for each bin using the 10 control urine samples, by picking one sample and calculating its AFE ratio using the mean AFE of the remaining 9 samples. An example for one healthy sample is shown in Equation 2. We repeated this for all 10 controls.

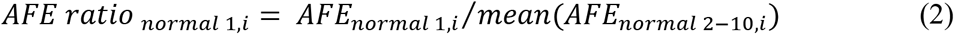

We then calculated the z-score of AFE ratio in bin *i* as the AFE ratio in bin *i* of a patient sample subtracted by the mean and divided by the standard deviation of background AFE ratio in bin *i* of the 10 healthy samples, as shown in Equation 3.

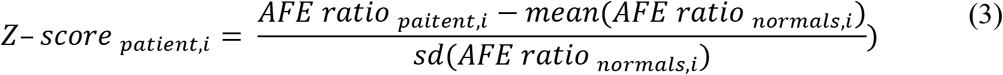

For each patient we compared the distribution of AFE ratio z-scores in copy number neural, loss, and gain bins.

### Urine Histone Analysis by Mass Spectrometry

As urine proteins are especially subject to hydrolysis due to urea, we isolated proteins encapsulated in extracellular vesicles (EVs) from urine to increase protein coverage. We isolated EVs from 10 ml pooled commercially available normal human urine (Lee Biosolutions, Maryland Heights, MO) using the ExoEasy Maxi kit (Qiagen, Germantown, MD) and following the manufacturer’s instructions. 4 mL of the flow-through fraction was processed by trichloroacetic acid (TCA) precipitation in 4:1 urine:acid ratio. Briefly, 1 ml of pre-chilled 100% TCA was added to 4 ml of urine flow-through, vortexed, and chilled for 1 h on ice. The sample was then centrifuged at 11,000 rcf for 30 min. After discarding the supernatant, pellets were first covered with 0.1% HCl in 100% ice cold acetone, then centrifuged at 11,000 rcf for 2 min. This step was repeated once again with 100% ice cold acetone. Pellets were then dried using nitrogen air flow and re-suspended in 200 μl of 50 mM ammonium bicarbonate for bicinchoninic acid (BCA) quantification. Equimolar amounts of the captured EV fraction and the TCA precipitated flow-through fraction were then diluted 2x with solution containing 50 mM Tris-HCl pH 7.0, 1X HALT (Thermo Fisher Scientific, San Jose, CA) and lysed by sonication on a UTR200 cup sonicator (Hielscher Ultrasound Technology, Teltow, Germany). Lysed fractions were incubated with TCEP (Thermo Fisher Scientific, Waltham, MA) at a final concentration of 5 mM for 45 min at 60 °C on thermoshaker, 450 rpm, followed by incubation with iodoacetemide (Sigma-Aldrich, Saint Louis, MO) to final concentration of 10 mM for 30 min at room temperature in the dark. Each fraction was then diluted three fold with 50 mM Tris-HCl. Polypeptides were trypsin digested at a ratio of 1:50 (Promega), overnight at 37 °C, and subjected to solid phase extraction. Peptides in solution were dried by speed vacuum and reconstituted in 50 mM NH_4_OH and quantified by BCA (Thermo Fisher Scientific). Basic reverse phase fractionation was carried out on 8 μg of tryptic peptides using an XBridge BEH C18 column (130 Å, 3.5 μm particle size, 4.6 mm x 100 mm) (Waters, Milford, MA) connected to a U3000 UHPLC (Thermo Fisher Scientific) system operating at 0.3 ml/min flow-rate. Peptides were fraction-collected into a 96-deep well plate using a gradient of acetonitrile and water, and 10% aqueous 50 mM Ammonium Hydroxide (pH 10)^27^. The resulting 96 fractions were concatenated into 6 analytical fractions, vacuum-dried and reconstituted in 6 μl of aqueous 0.1% formic acid solution for LC-MS/MS analysis.

Mass spectrometry acquisition was performed in top-speed data-dependent mode (3 second duty cycle) on an Orbitrap Fusion Lumos Tribrid (Thermo Fisher Scientific) mass spectrometer coupled to a nanoAcquity UPLC system (Waters). Peptides were separated on a PepMap RSLC C18 EasySpray C18 column (100 Å, 2 μm particle size, 75 μm x 25 cm) kept at 50 °C with a 120 min gradient from 3% to 30% to 90% acetonitrile in 0.1% formic acid, at a flow-rate of 350 nl/min. The mass spectrometer was operated with the following parameters: ion transfer tube temperature of 275 °C, spray voltage of 2400 V, MS1 in Orbitrap with a resolution of 120K and mass range of 400-1500 m/z, most abundant precursors (excluding undetermined and +1 charge state species) were selected for MS2 measurement in the iontrap following HCD fragmentation with 35% collision energy; dynamic exclusion was set to 60 s. Mass spectra were searched using Proteome Discoverer (v2.1.0.388, Thermo Fisher Scientific) and Mascot (Matrix Science, Boston, MA) on a human UniprotKB (Swissprot, June 2017) database allowing for two missed cleavages, fixed cysteine carbamidomethylation and variable methionine oxidation, a 10 ppm precursor and 0.6 Da fragment mass tolerance. Percolator was employed with a target-decoy strategy to determine false discovery rates at peptide and protein level^28^.

## Supplementary Materials

Supplementary Figure 1: Fragment size distributions in individual control plasma samples.

Supplementary Figure 2: Fragment size distributions in individual control urine samples.

Supplementary Figure 3: Comparison of raw sequencing coverage between plasma and urine.

Supplementary Figure 4: Nucleotide frequencies at fragment start sites in plasma samples.

Supplementary Figure 5: Nucleotide frequencies at fragment end sites in plasma samples.

Supplementary Figure 6: Nucleotide frequencies at fragment start sites in urine samples.

Supplementary Figure 7: Nucleotide frequencies at fragment end sites in urine samples.

Supplementary Figure 8: Nucleotide frequencies at fragment start and end sites across fragment size bins.

Supplementary Figure 9: Comparison of overlapping and non-overlapping nucleosome call confidence scores.

Supplementary Figure 10: Distance of fragment start and end sites relative to nucleosome dyad using a urine-based nucleosome occupancy map.

Supplementary Figure 11: Cosine similarity between cfDNA fragment sizes and DHS sites.

Supplementary Figure 12: Spearman’s rank correlation coefficients for NDR coverage and gene expression.

Supplementary Figure 13: Rank changes in correlation between NDR coverage and gene expression between plasma and urine.

Supplementary Figure 14: Fragment size distribution in individual urine samples from cancer patients.

Supplementary Figure 15: Nucleotide frequencies at fragment start sites in urine samples from cancer patients.

Supplementary Figure 16: Nucleotide frequencies at fragment end sites in urine samples form cancer patients.

Supplementary Figure 17: ROC analysis for classifying control and cancer samples by cancer type

Supplementary Figure 18: Aberrant fragment ends (AFE) across copy number changes in sample 50.

Supplementary Figure 19: Aberrant fragment ends (AFE) across copy number changes in sample 43.

Supplementary Figure 20: Aberrant fragment ends (AFE) across copy number changes in sample 36.

Supplementary Figure 21: Aberrant fragment ends (AFE) across copy number changes in sample 37.

Supplementary Figure 22: Aberrant fragment ends (AFE) across copy number changes in sample 34.

Supplementary Figure 23: Aberrant fragment ends (AFE) across copy number changes in sample 33.

Supplementary Table 1: Histone proteins identified in extracellular vesicles isolated from urine using mass spectrometry.

Supplementary Table 2: Clinical characteristics of cancer patients.

Supplementary Data 1: Quantile normalized cosine similarity between DHS sites and cfDNA fragment size.

Supplementary Data 2: Quantile normalized Spearman’s rho between gene expression and NDR coverage.

Supplementary Data 3: Rank changes in Spearman’s rho across pooled plasma and urine.

Supplementary Data 4: Aberrant fragment ends (AFE) and multidimensional scaled dimensions 1-4 of aberrant fragment end motifs (AFEM) in urine samples.

## Acknowledgements

We would like to thank Callie Sinclair and Stephanie Buchholtz at TGen, Lisa Keller at Phoenix Children’s Hospital, volunteers and patients who participated in this study.

## Author Contributions

HM and MM conceptualized and designed the study. HM, JZ, TCC, ER, AO, PP, KV and MM developed methods. JZ, MCM, CB, SAC, AG, DDV and PH designed and conducted prospective clinical studies. TCC, ER, AO, SC, EH and MMG generated data. HM, EH and BRM analyzed sequencing data. HM, ER, PP and MM interpreted data. HM and MM wrote the paper with assistance from JZ, TCC, ER, DVH, SAC, PH and PP. All authors approved the final manuscript.

## Data and materials availability

Urine and plasma sequencing data from controls will be made available in dbGaP upon manuscript acceptance. Urine sequencing data and tumor/germline exome sequencing data from pediatric cancer patients will be made available in dbGaP upon manuscript acceptance. Tumor/germline sequencing data and urine sequencing data from pancreatic cancer patients will be made available upon reasonable request to authors.

