## Supplementary Materials for "Sub-nucleosomal organization in urine cell-free DNA"

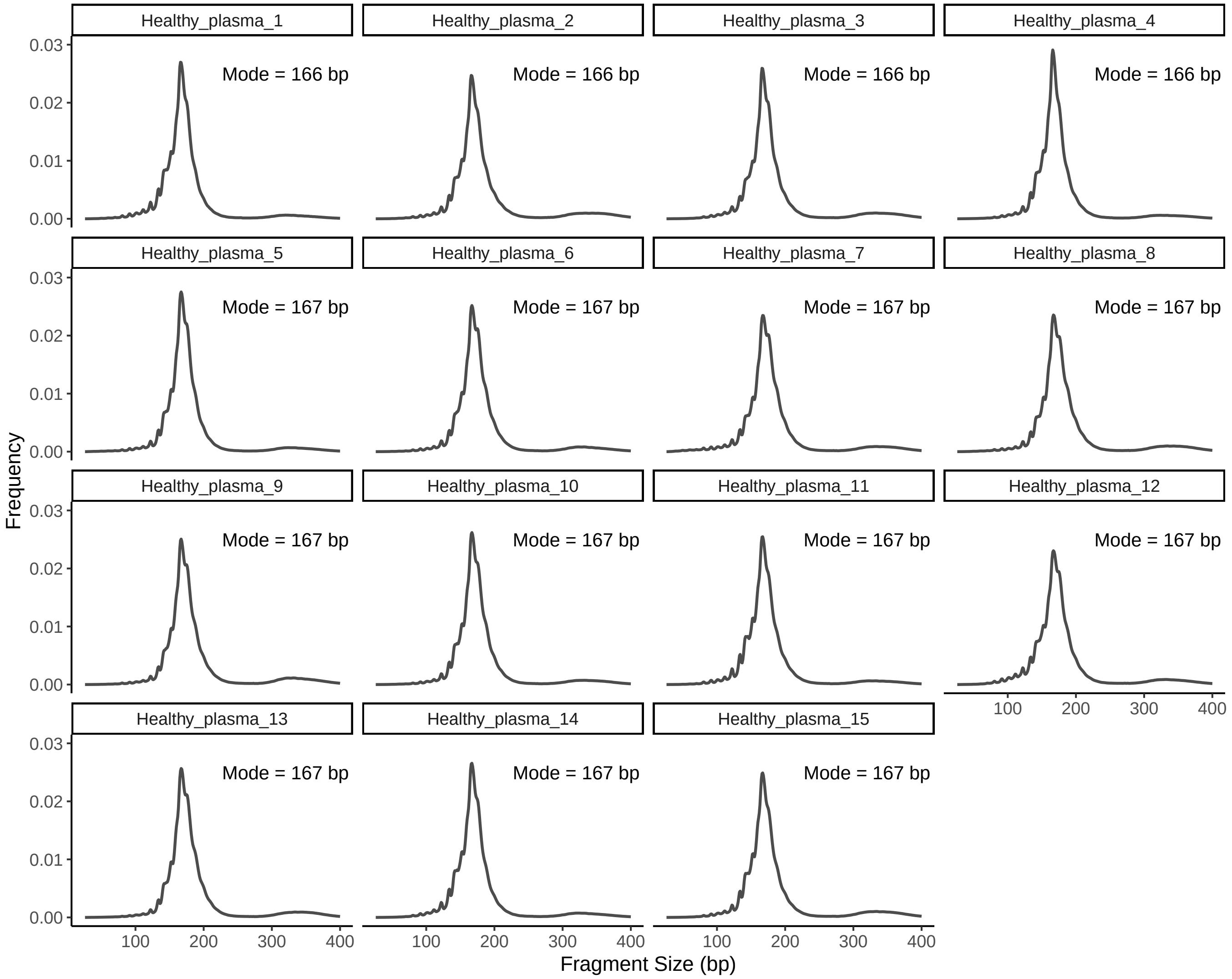


**Supplementary Figure 1: Fragment size distributions in individual control plasma samples.**

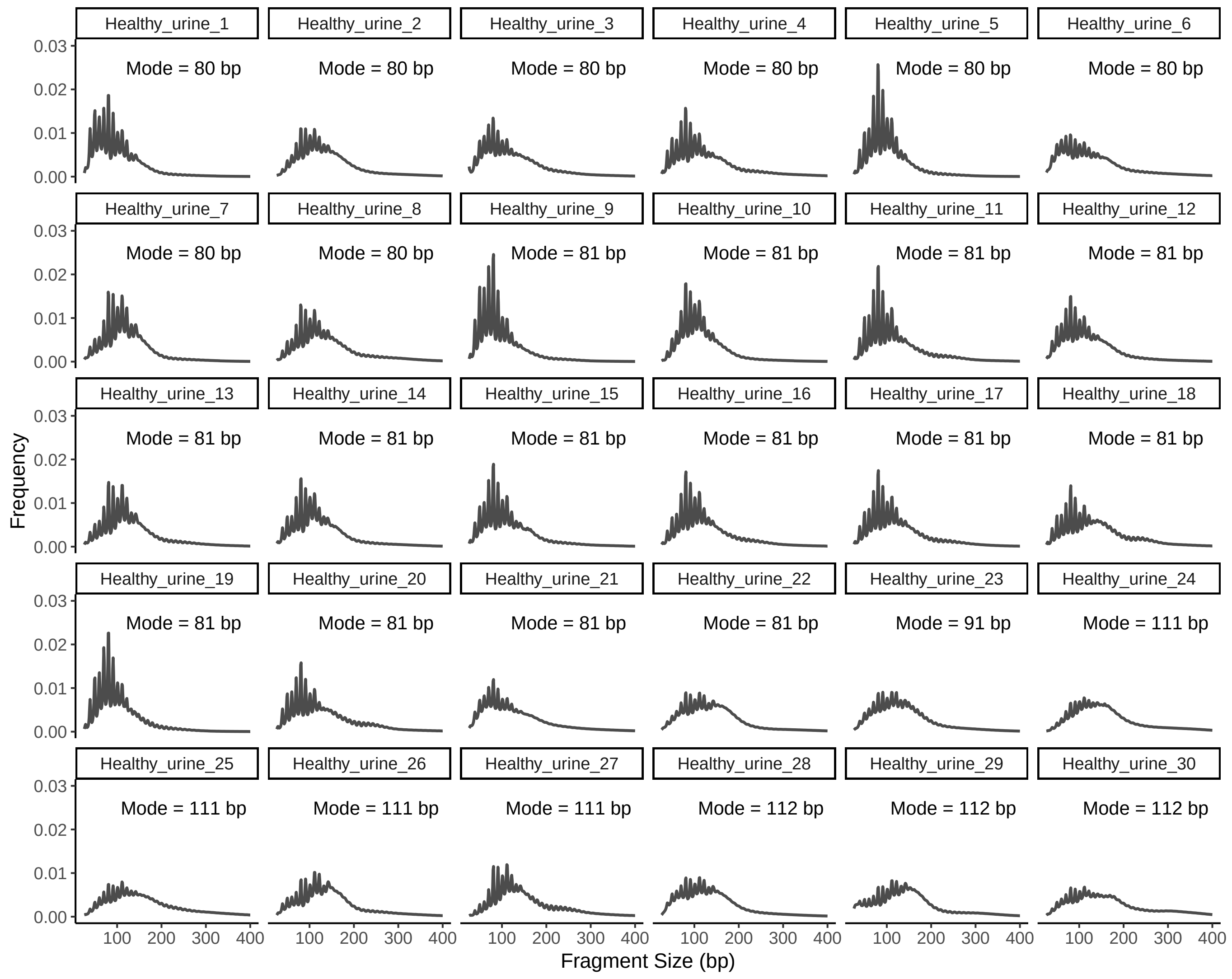


**Supplementary Figure 2: Fragment size distributions in individual control urine samples.**

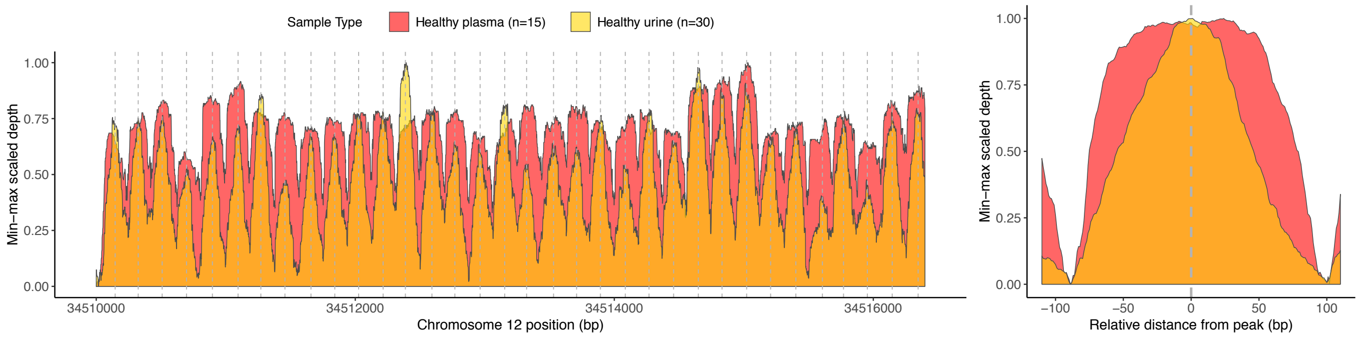


**Supplementary Figure 3: Comparison of raw sequencing coverage between plasma and urine.** Raw min-max scaled physical sequencing coverage of pooled plasma and urine samples in a genomic region with stable nucleosomes (Chromosome 12p11.1).

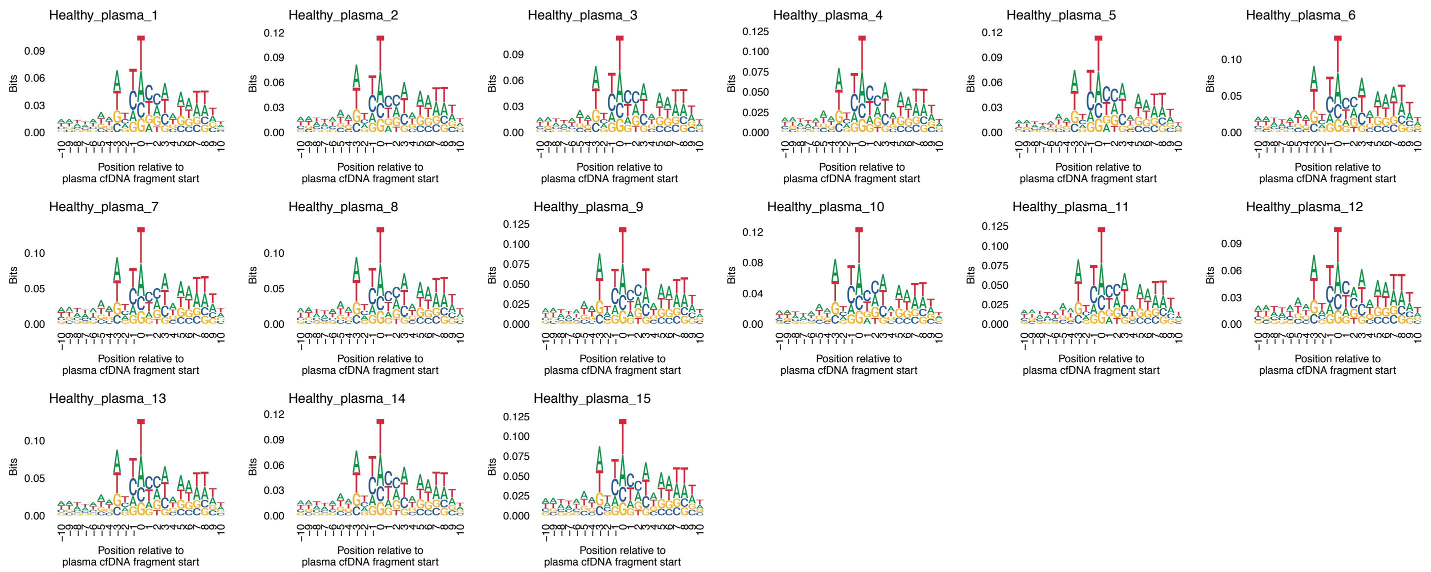


**Supplementary Figure 4: Nucleotide frequencies at fragment start sites in plasma samples.**

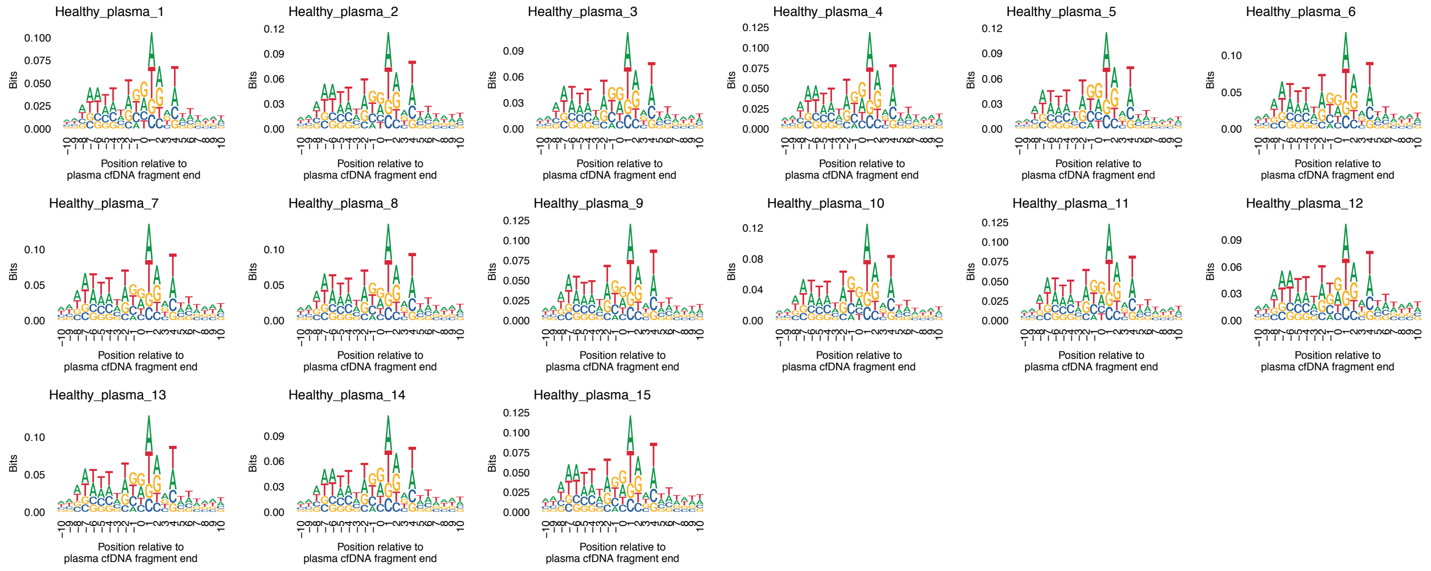


**Supplementary Figure 5: Nucleotide frequencies at fragment end sites in plasma samples.**

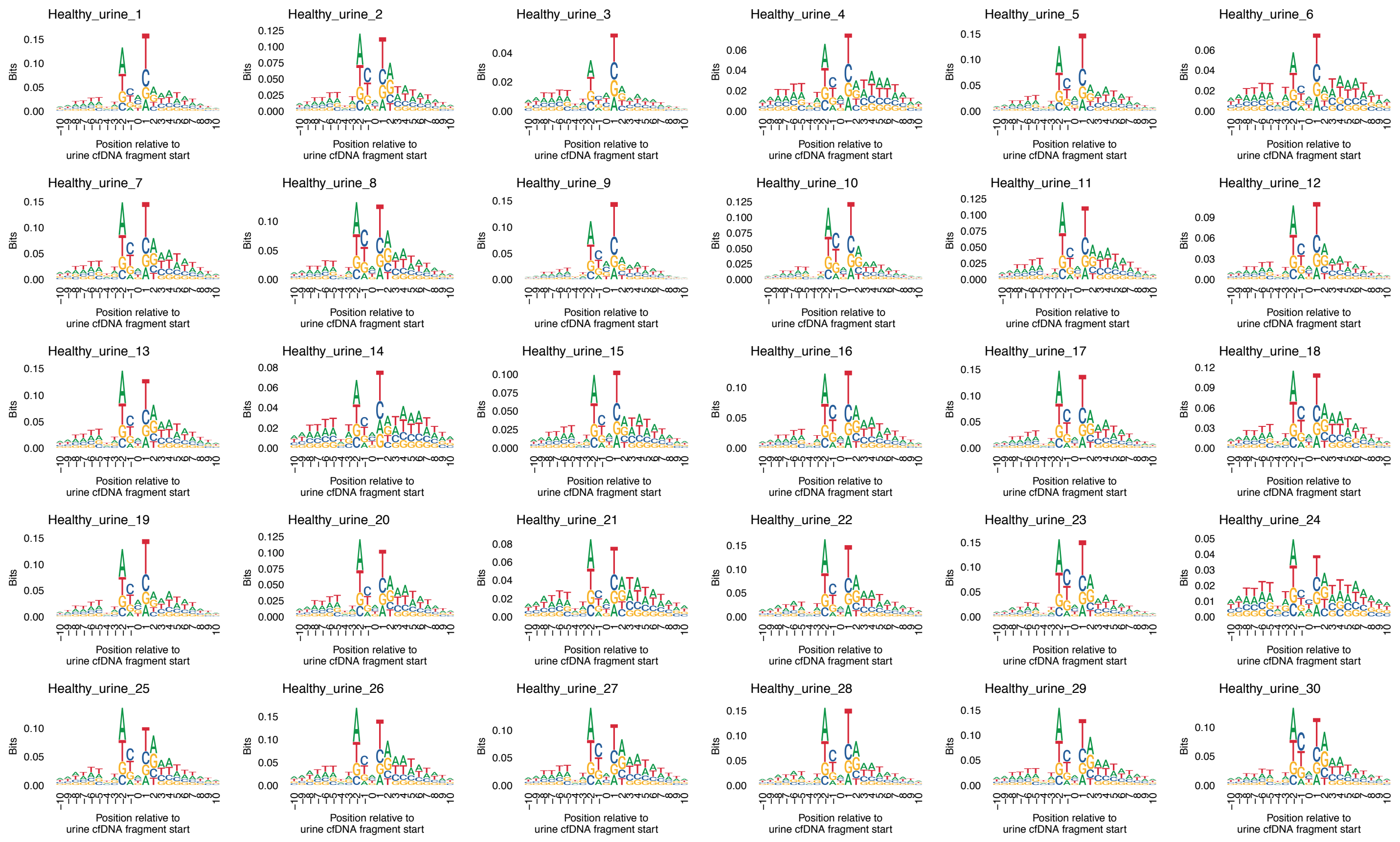


**Supplementary Figure 6: Nucleotide frequencies at fragment start sites in urine samples.**

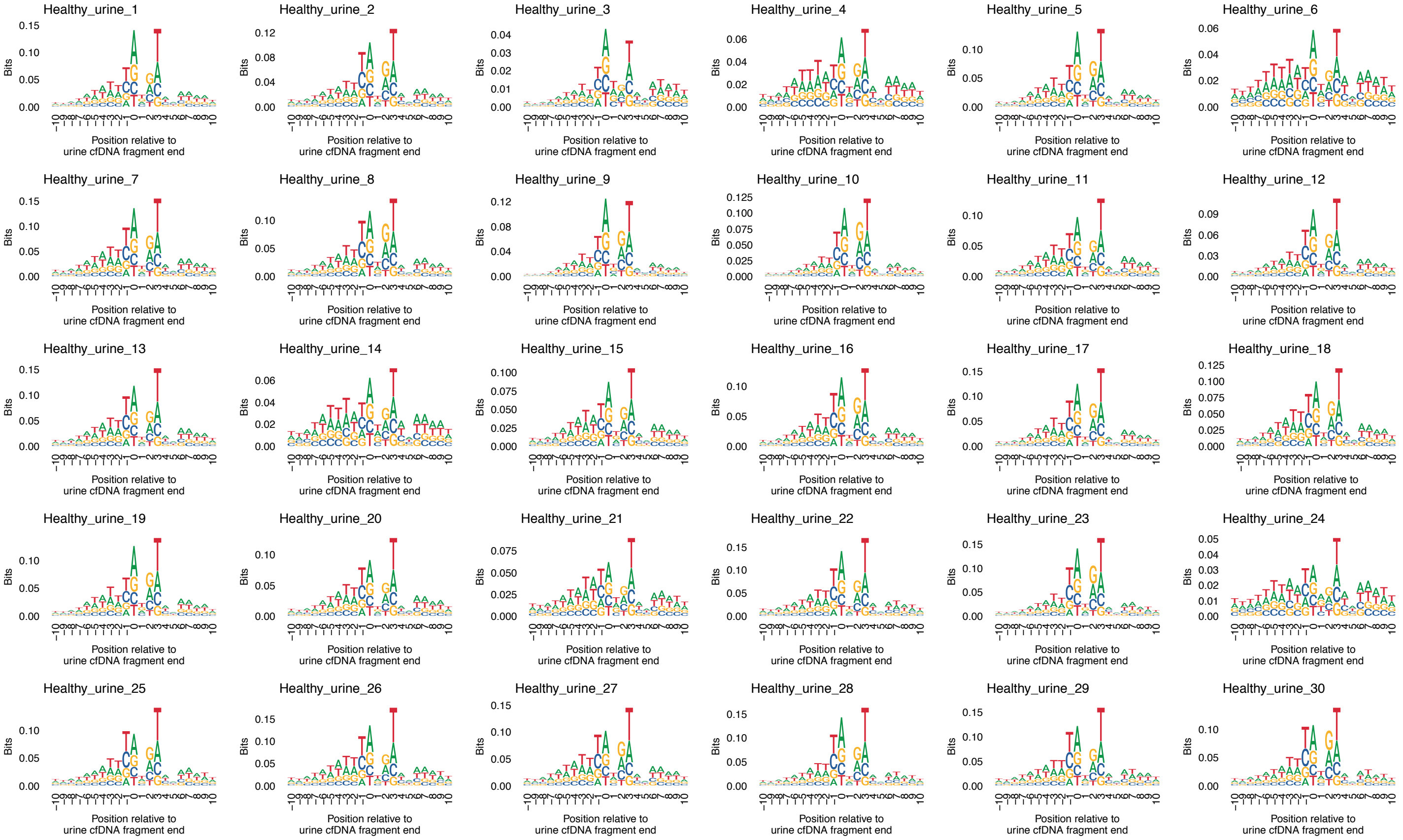


**Supplementary Figure 7: Nucleotide frequencies at fragment end sites in urine samples.**

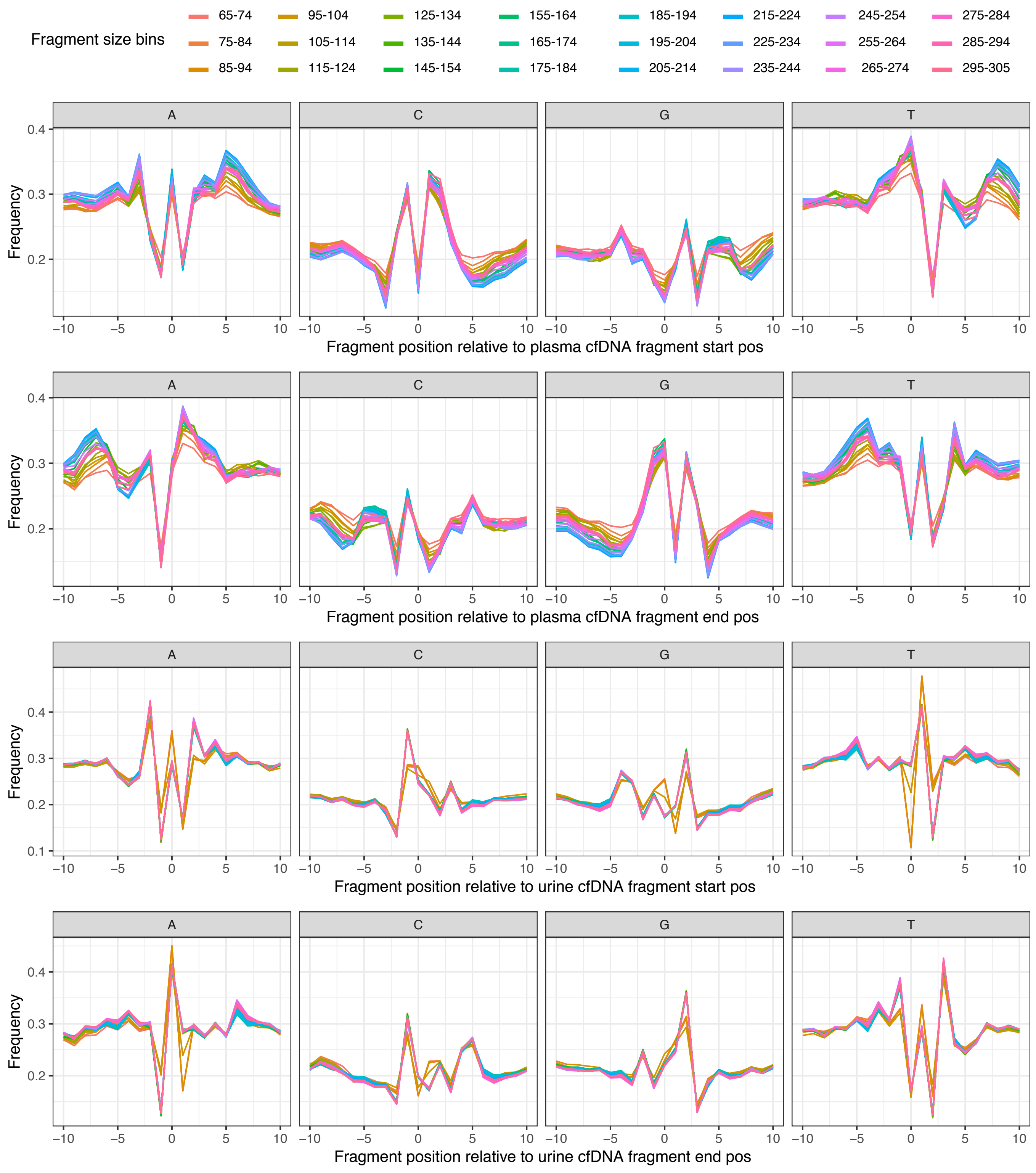


**Supplementary Figure 8: Nucleotide frequencies at fragment start and end sites across fragment size bins.**

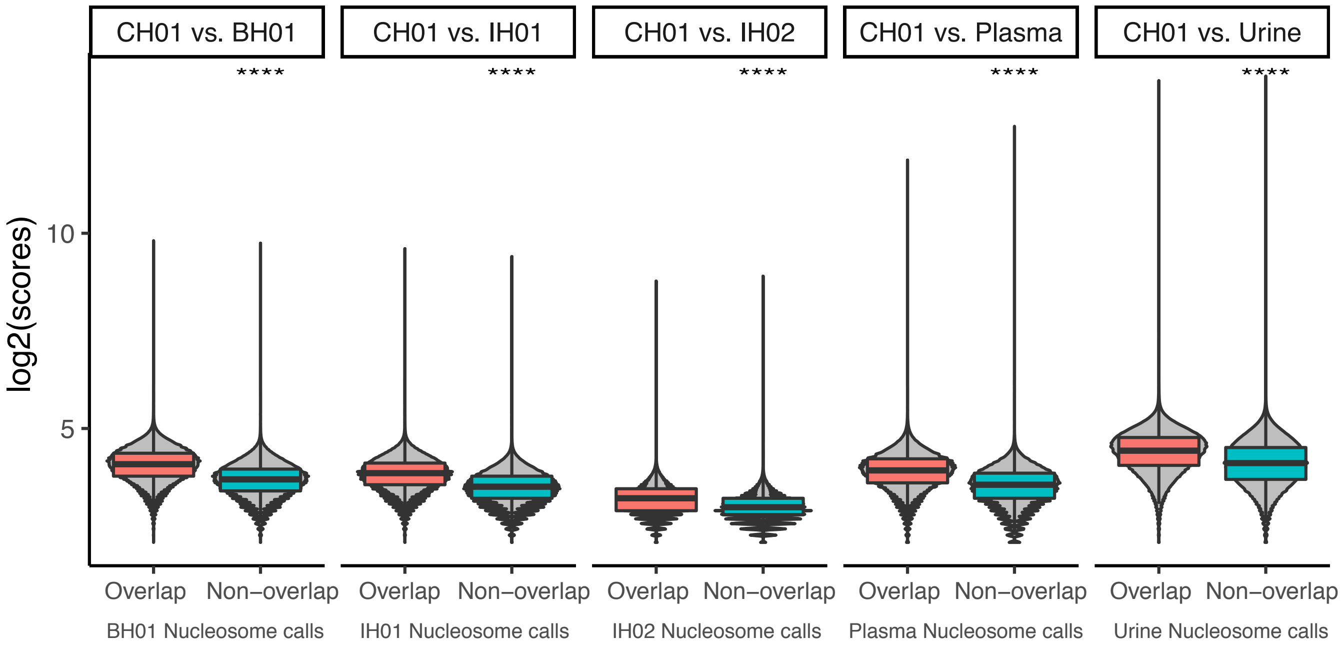


**Supplementary Figure 9: Comparison of overlapping and non-overlapping nucleosome call confidence scores.** Pooled urine, pooled plasma, IH01, IH02, and BH01 nucleosome tracks were compared with CH01 nucleosome track. The **** represents t.test two-tailed p-value $< 0.00001$.

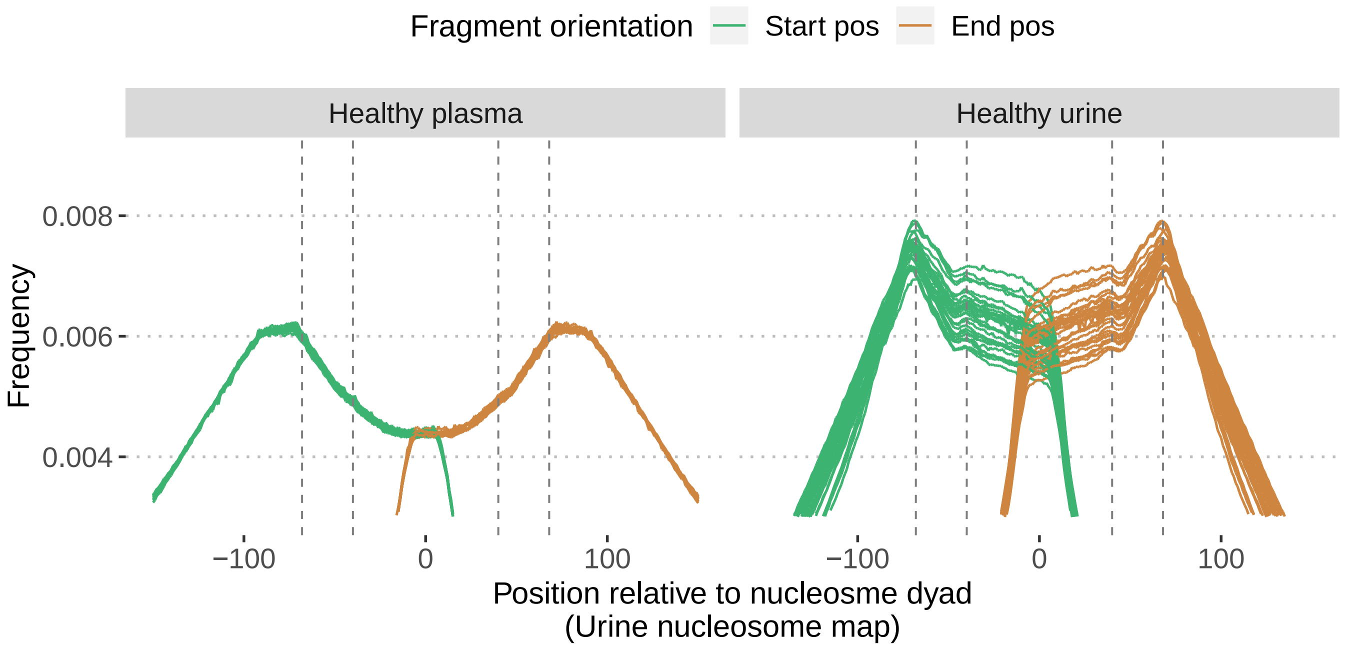


**Supplementary Figure 10: Distance of fragment start and end sites relative to nucleosome dyad using a urine-based nucleosome occupancy map.** The vertical lines are drawn at 40 bp and 68 bp downstream and upstream from the nucleosome dyad.

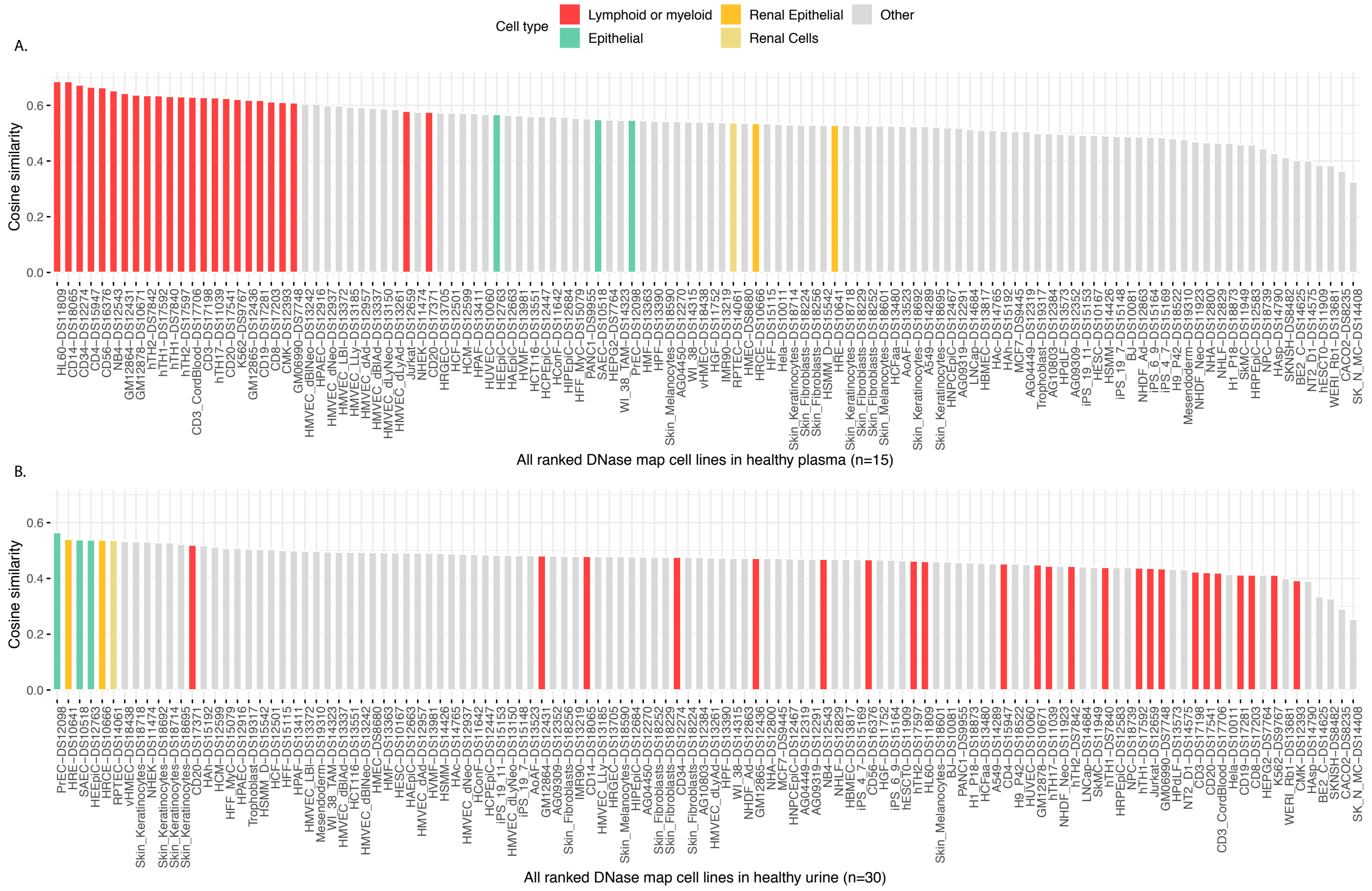


**Supplementary Figure 11: Cosine similarity between cfDNA fragment sizes and DHS sites.** (A) Cosine similarities were calculated between pooled control plasma sample z-score normalized median cfDNA fragment sizes and modified z-score normalized number of DNase hypersensitivity sites in 500kb bins. (B) Same analysis as (A) except using pooled control urine sample.

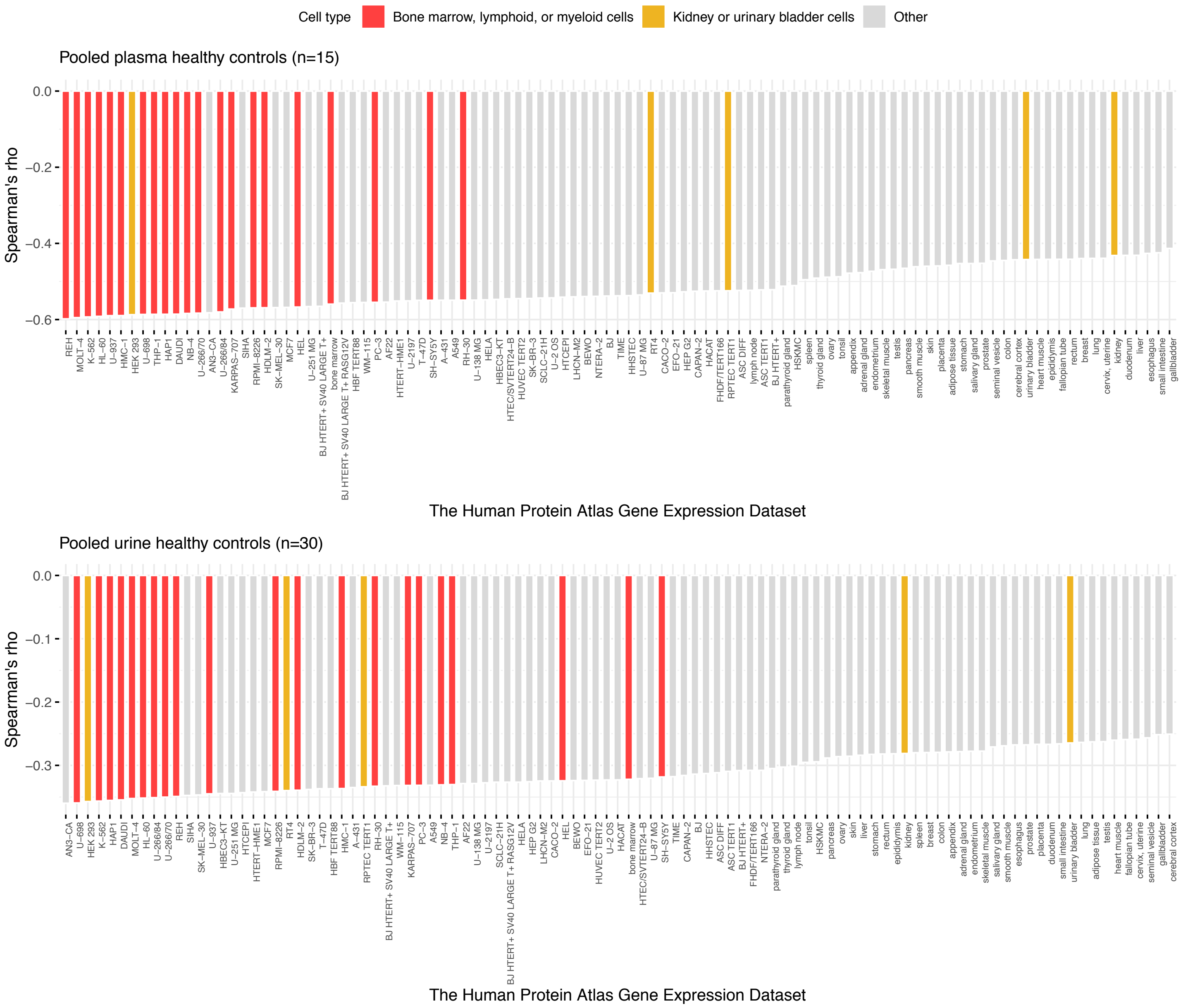


**Supplementary Figure 12: Spearmans rank correlation coefficients for nucleosome depleted region (NDR) coverage and gene expression.** Gene expression data sets from 64 human cell lines and 37 primary tissues (Human Protein Atlas) ranked based on Spearman's rank correlation coefficient (Spearman's rho) between mean TSS coverage of pooled plasma (A) and urine (B) samples and gene expression values from individual data sets.

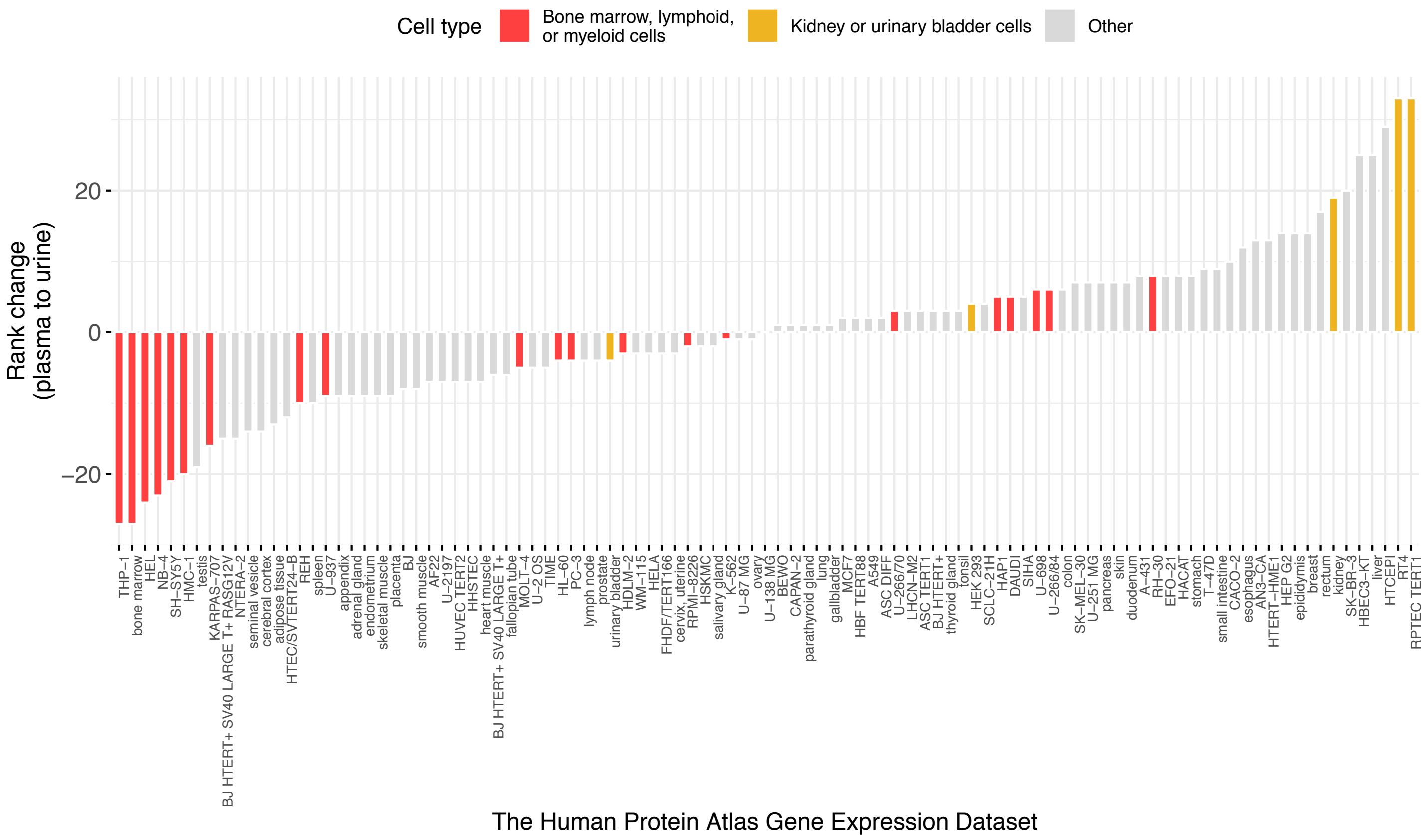


**Supplementary Figure 13: Rank changes in correlation between NDR coverage and gene expression between plasma and urine.**

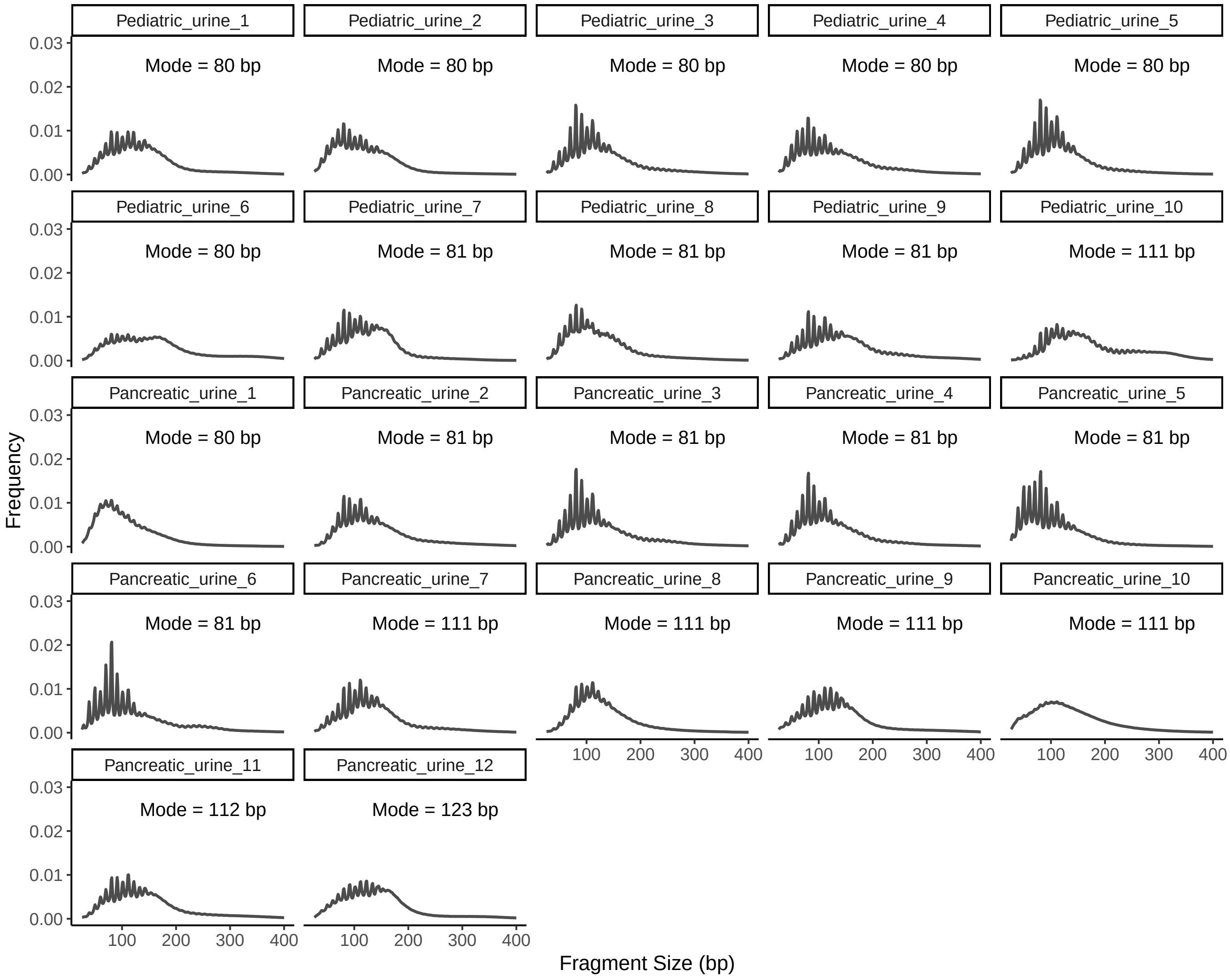


**Supplementary Figure 14: Fragment size distribution in individual urine samples from cancer patients.**

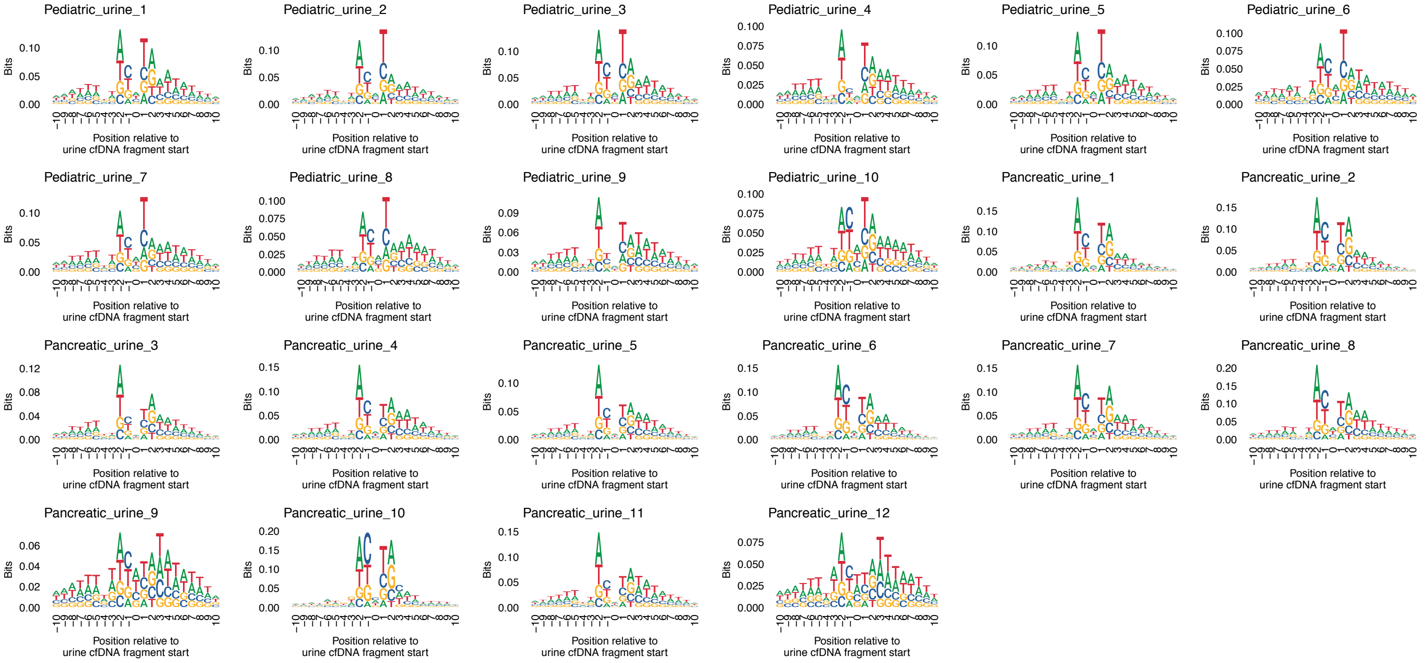


**Supplementary Figure 15: Nucleotide frequencies at fragment start sites in urine samples from cancer patients.**

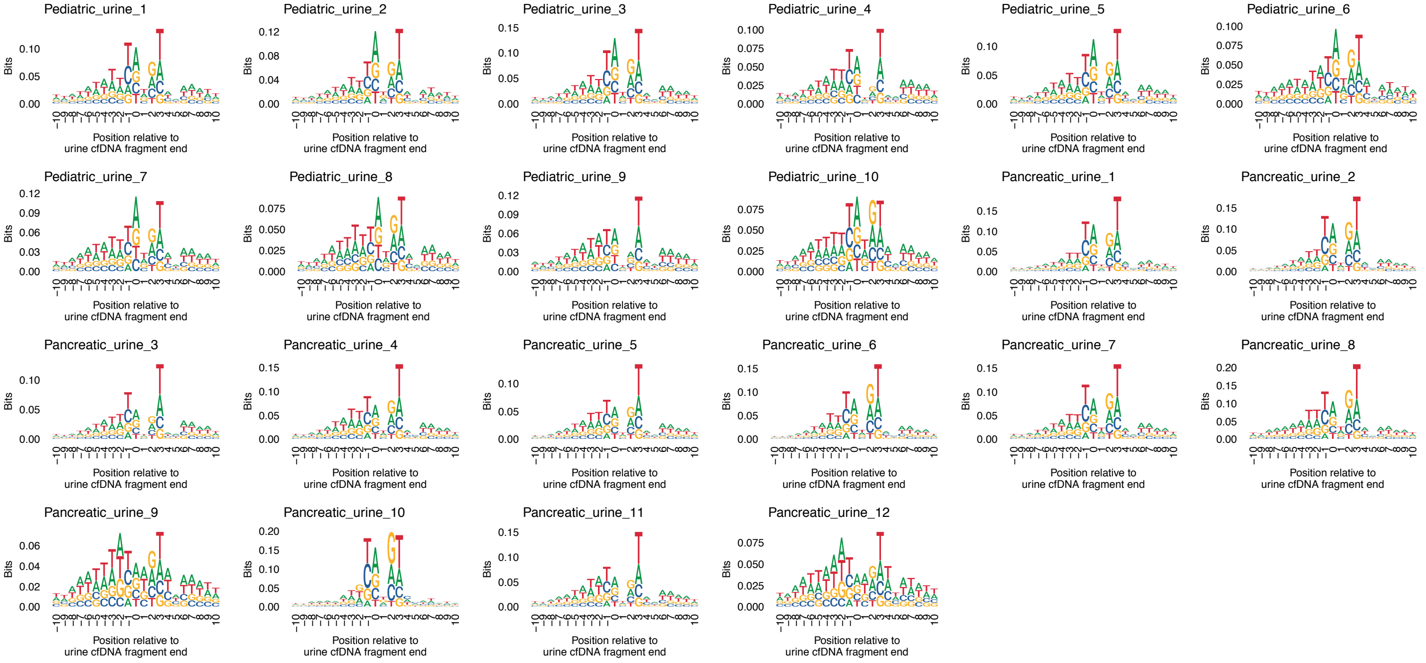


**Supplementary Figure 16: Nucleotide frequencies at fragment end sites in urine samples form cancer patients.**

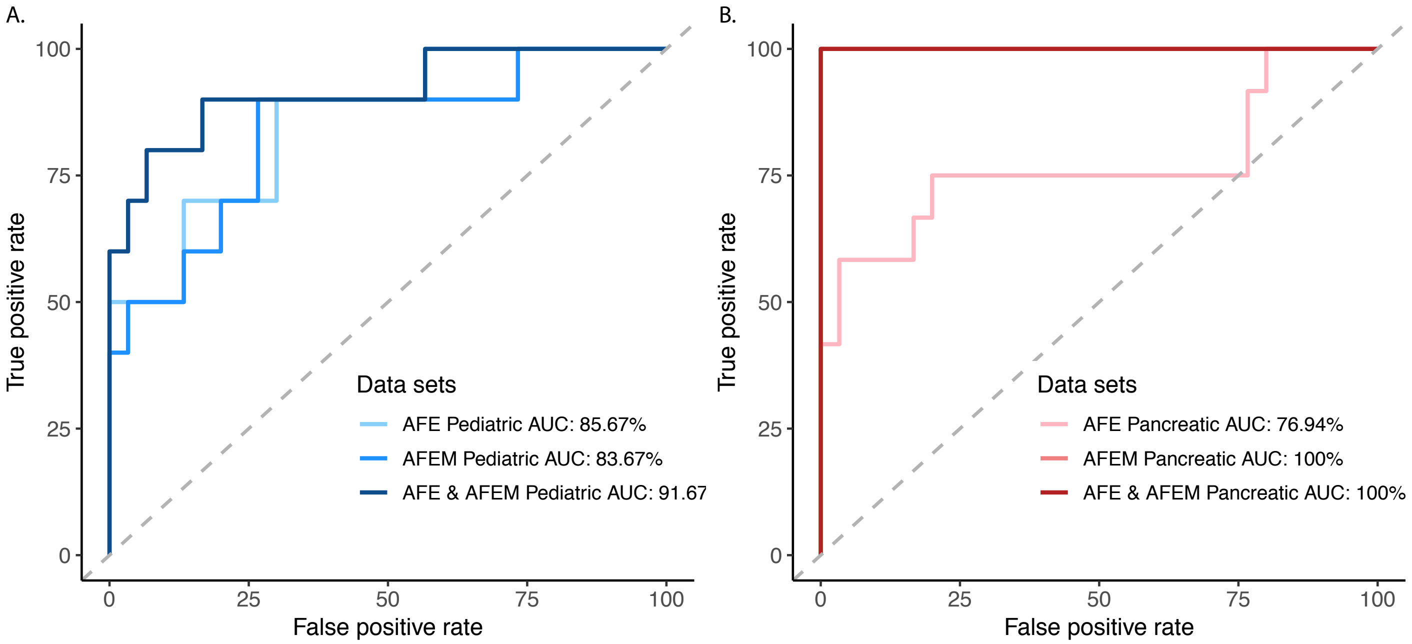


**Supplementary Figure 17: ROC analysis for classifying control and cancer samples by cancer type.** For AFE analysis, the fractions shown in Figure 6B were used for ROC analysis. For AFEM and combined AFE and AFEM, probabilities from a logistic regression fitted to the first 4 MDS dimensions and AFE were used for ROC analysis. (A-B) ROC analysis for pediatric and pancreatic cancer patients respectively.

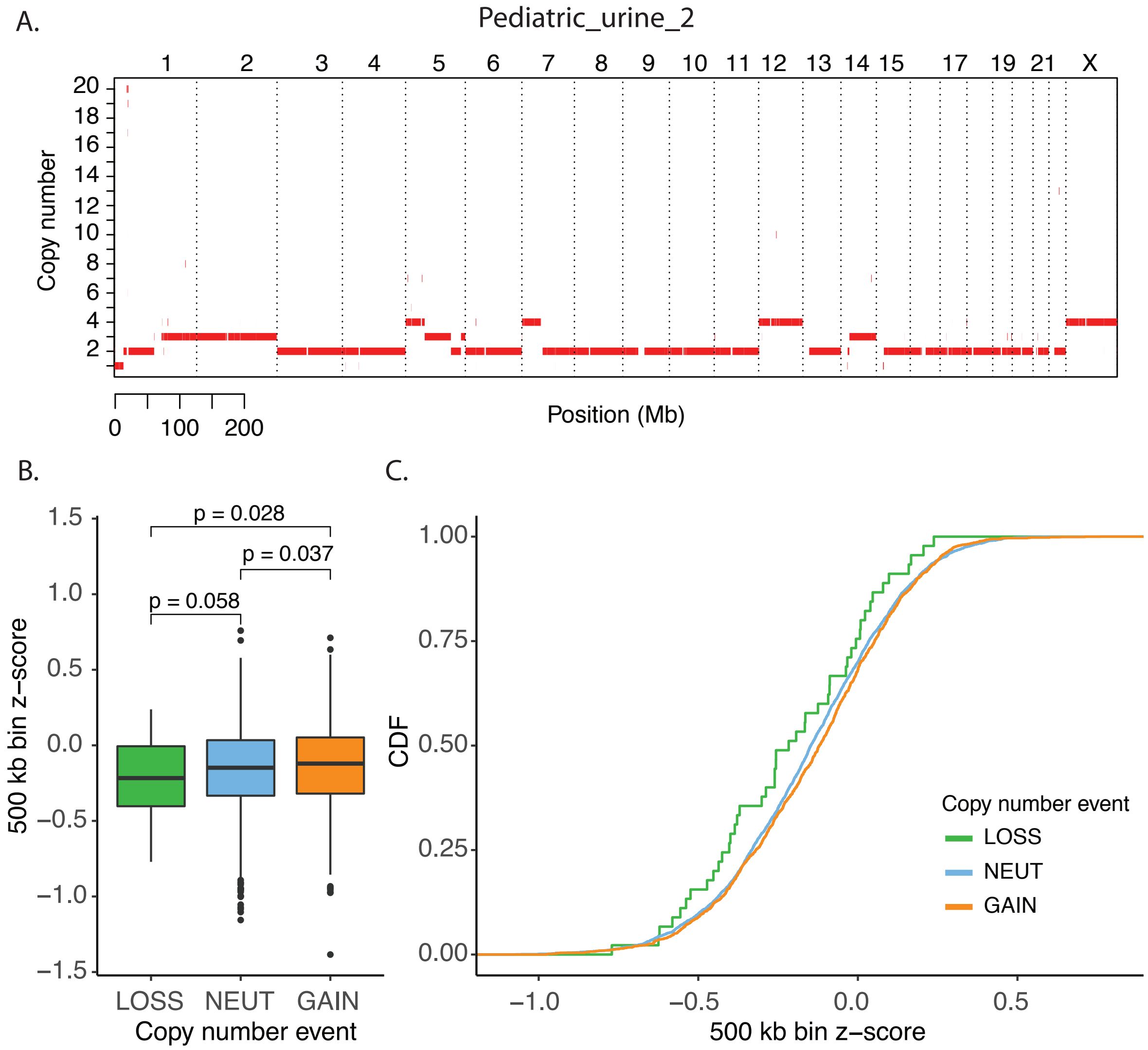


**Supplementary Figure 18: Aberrant fragment ends (AFE) across copy number changes in sample 50.** (A) Copy number aberration analysis from tumor and normal sequencing of sample 50 (Pediatric_urine_2). (B) 500kb bins where copy number was 2, $< 2$, and $> 2$ was labeled as neutral, loss, and gain. AFE in 500kb bins were converted to a z-score based on a background distribution of per bin AFE from healthy control samples. The z-score was compared between neutral, loss, and gain bins using a one-tailed t.test. (C) The cumulative distribution of AFE in 500kb bin z-score in neutral, loss, and gain bins.

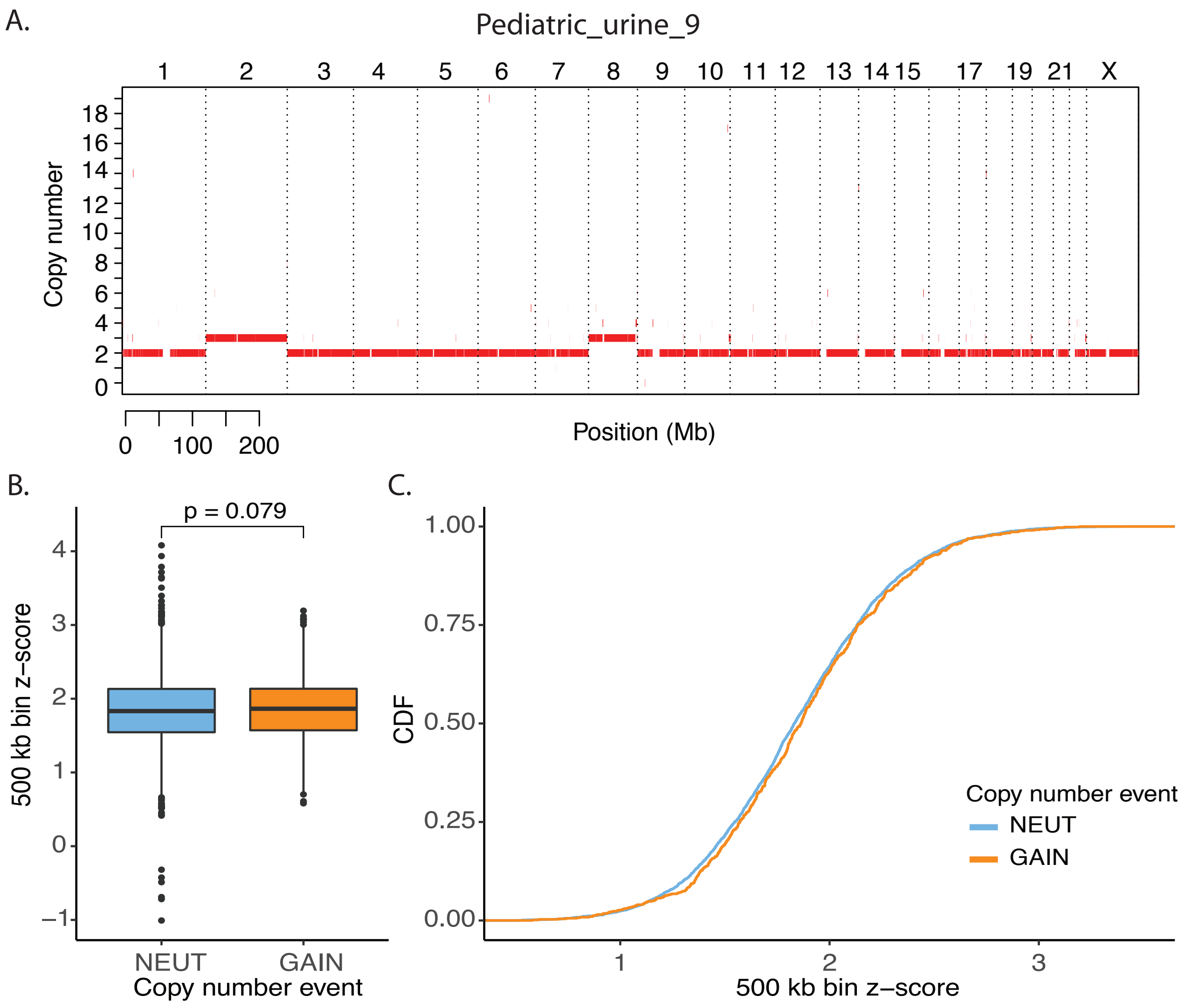


**Supplementary Figure 19: Aberrant fragment ends (AFE) across copy number changes in sample 43.** (A) Copy number aberration analysis from tumor and normal sequencing of sample 43 (Pediatric_urine_9). (B) 500 kb bins where copy number was 2, $< 2$, and $> 2$ was labeled as neutral, loss, and gain. AFE in 500 kb bins were converted to a z-score based on a background distribution of per bin AFE from healthy control samples. The z-score was compared between neutral, loss, and gain bins using a one-tailed t.test. (C) The cumulative distribution of AFE in 500 kb bin z-score in neutral, loss, and gain bins. In this patient there were not enough bins with copy number $< 2$, thus loss bins are not shown.

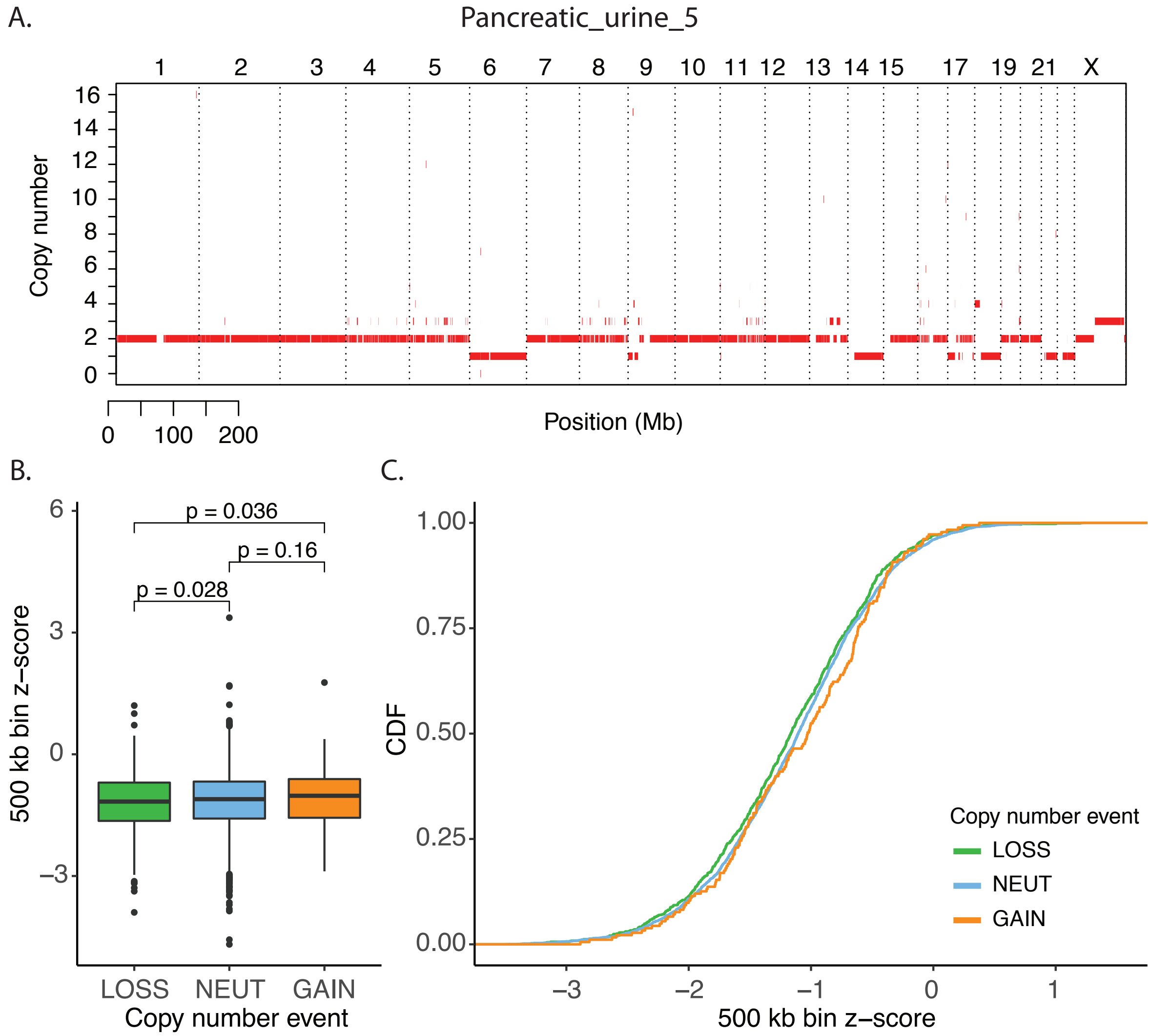


**Supplementary Figure 20: Aberrant fragment ends (AFE) across copy number changes in sample 36.** (A) Copy number aberration analysis from tumor and normal sequencing of sample 36 (Pancreatic_urine_5). (B) 500 kb bins where copy number was 2, $< 2$, and $> 2$ was labeled as neutral, loss, and gain. AFE in 500 kb bins were converted to a z-score based on a background distribution of per bin AFE from healthy control samples. The z-score was compared between neutral, loss, and gain bins using a one-tailed t.test. (C) The cumulative distribution of AFE in 500 kb bin z-score in neutral, loss, and gain bins.

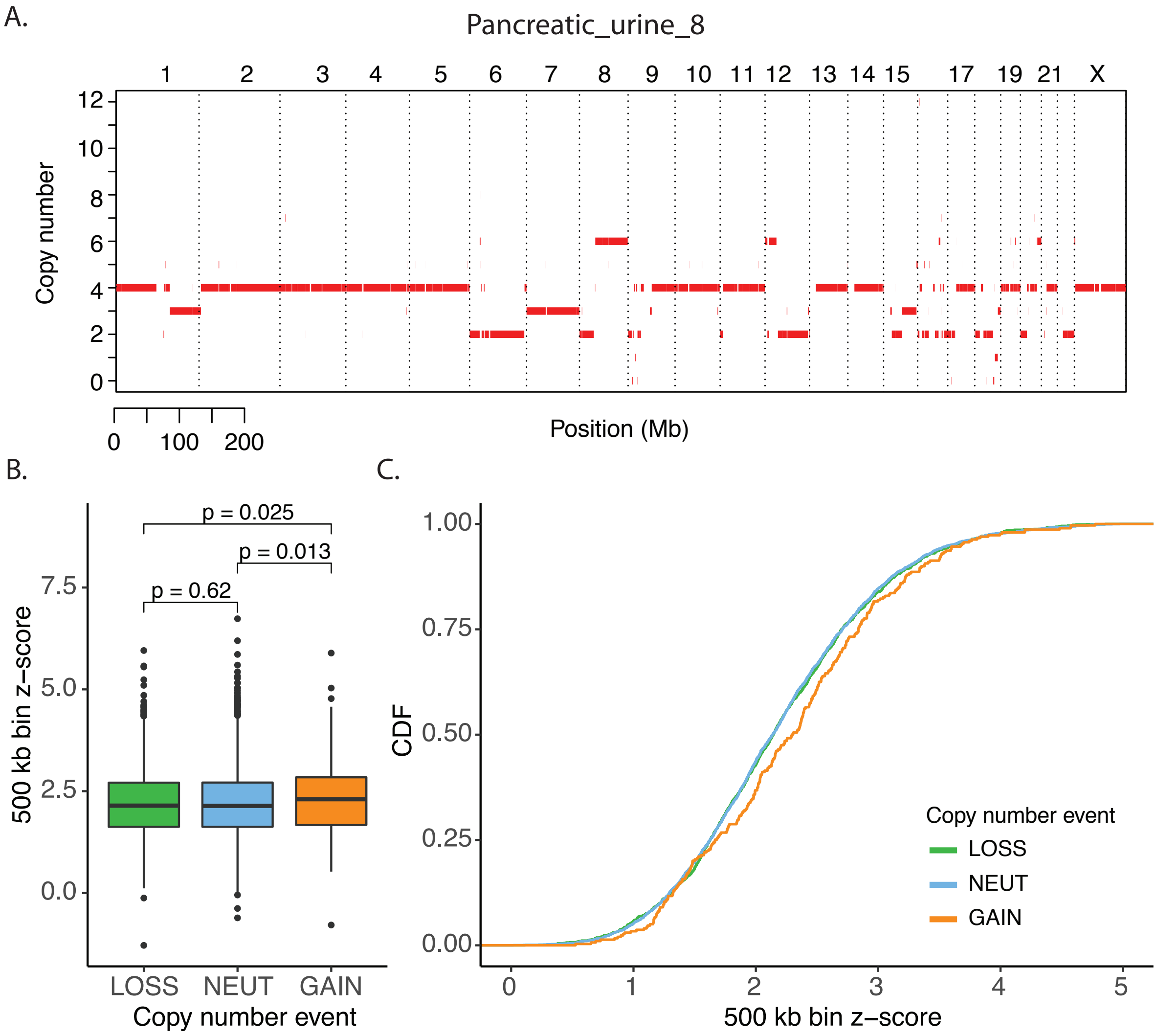


**Supplementary Figure 21: Aberrant fragment ends (AFE) across copy number changes in sample 37.** (A) Copy number aberration analysis from tumor and normal sequencing of sample 37 (Pancreatic_urine_8). (B) Since the patient seemed to have a genome wide duplication event, 500 kb bins where copy number was 4, $< 4$, and $> 4$ was labeled as neutral, loss, and gain. AFE in 500 kb bins were converted to a z-score based on a background distribution of per bin AFE from healthy control samples. The z-score was compared between neutral, loss, and gain bins using a one-tailed t.test. (C) The cumulative distribution of AFE in 500 kb bin z-score in neutral, loss, and gain bins.

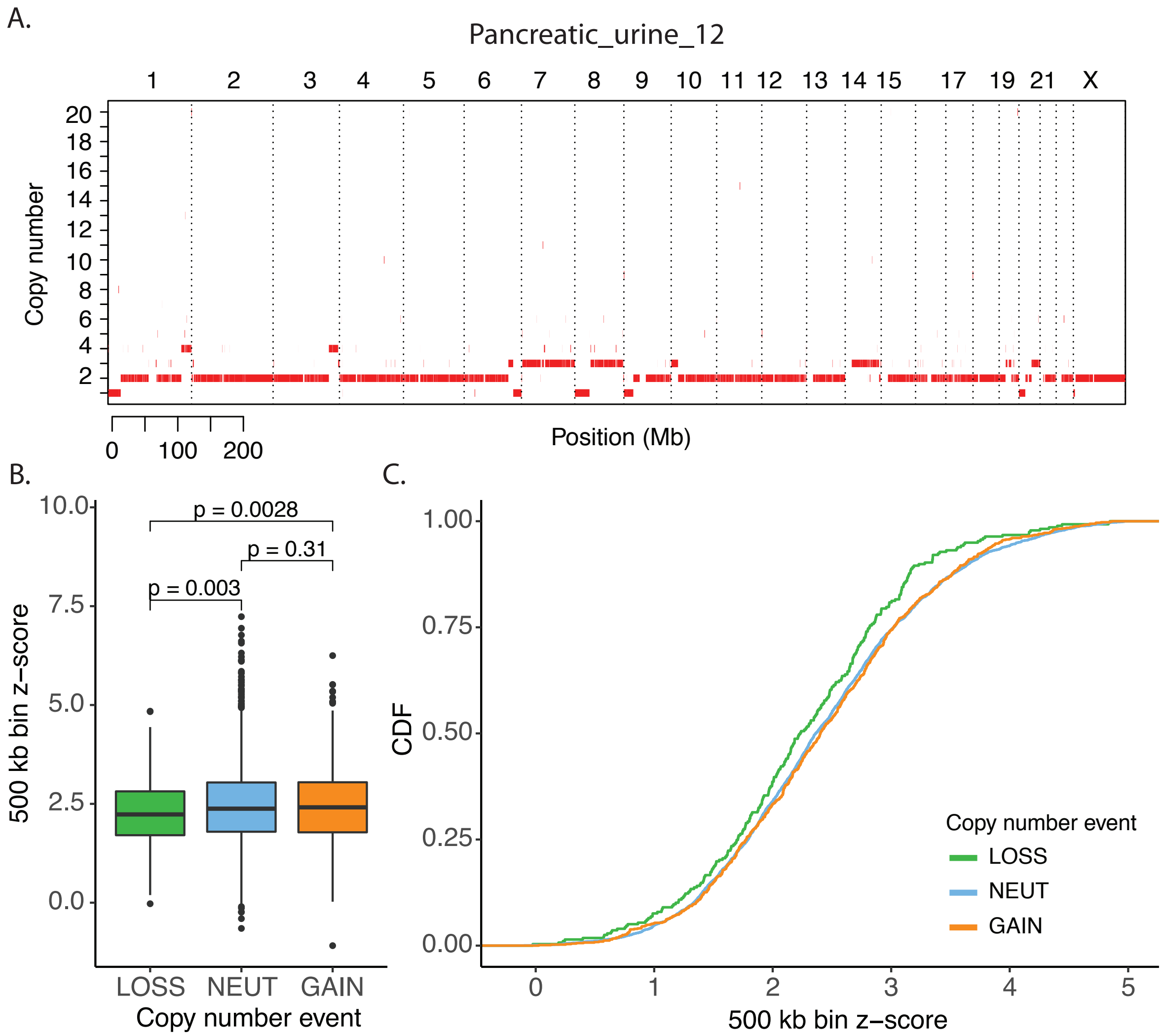


**Supplementary Figure 22: Aberrant fragment ends (AFE) across copy number changes in sample 34.** (A) Copy number aberration analysis from tumor and normal sequencing of sample 34 (Pancreatic_urine_12). (B) 500 kb bins where copy number was 2, $< 2$, and $> 2$ was labeled as neutral, loss, and gain. AFE in 500 kb bins were converted to a z-score based on a background distribution of per bin AFE from healthy control samples. The z-score was compared between neutral, loss, and gain bins using a one-tailed t.test. (C) The cumulative distribution of AFE in 500 kb bin z-score in neutral, loss, and gain bins.

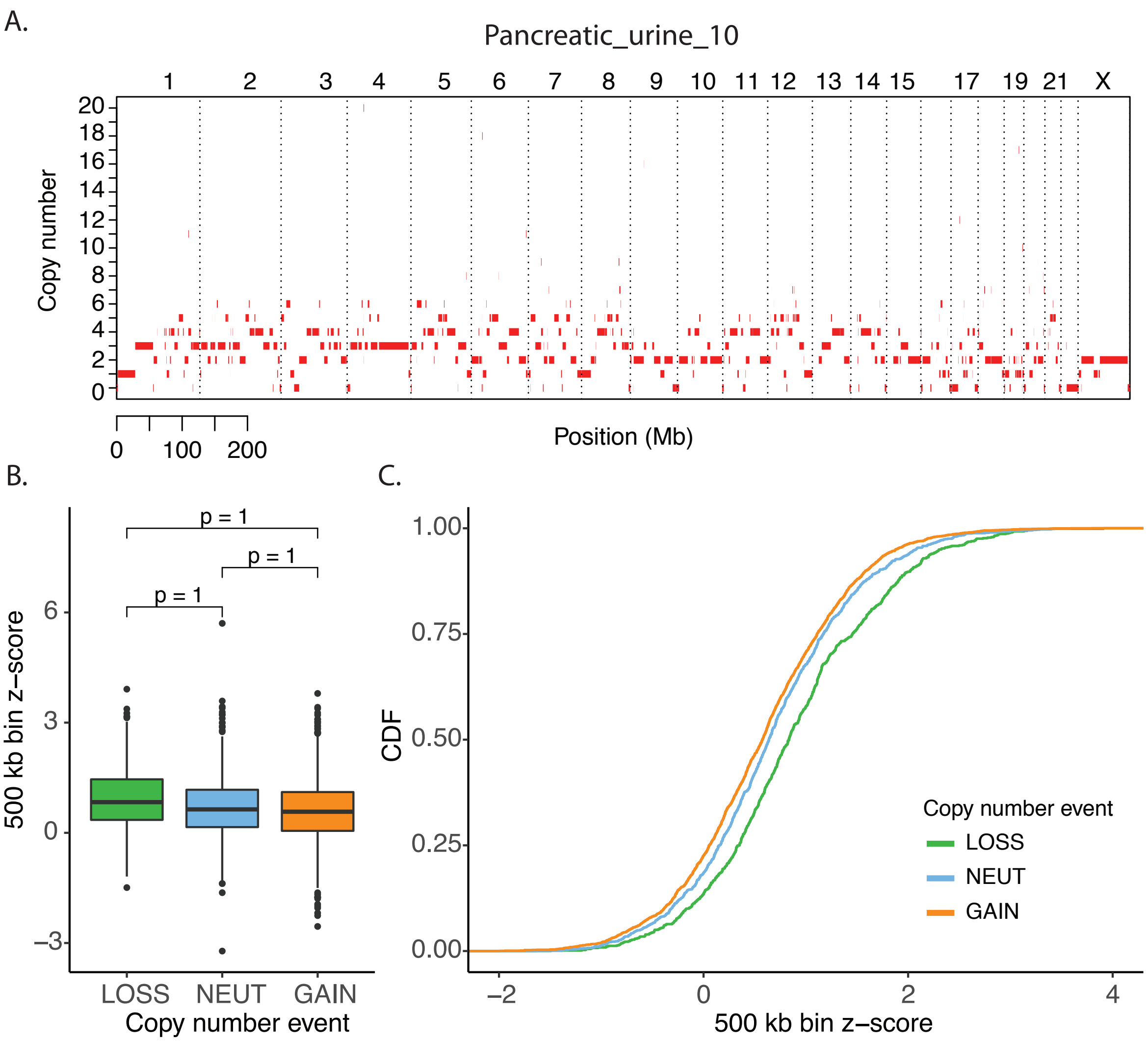


**Supplementary Figure 23: Aberrant fragment ends (AFE) across copy number changes in sample 33.** (A) Copy number aberration analysis from tumor and normal sequencing of sample 33 (Pancreatic_urine_10). (B) 500 kb bins where copy number was 2, < 2, and > 2 was labeled as neutral, loss, and gain. AFE in 500 kb bins were converted to a z-score based on a background distribution of per bin AFE from healthy control samples. The z-score was compared between neutral, loss, and gain bins using a one-tailed t.test. (C) The cumulative distribution of AFE in 500 kb bin z-score in neutral, loss, and gain bins. We believe this patient shows an unexpected distribution of AFE because this might be a case of chromothripsis and consistent copy number regions are hard to infer.

| Gene Name | Accession | Description | Coverage [%] | Peptides [#] | PSMs [#] |
|---|---|---|---|---|---|
| H1F0 | P07305 | Histone H1.0 | 9 | 2 | 2 |
| HIST1H2AB | P04908 | Histone H2A type 1-B/E | 30 | 3 | 8 |
| HIST2H2BF | Q5QNW6-2 | Isoform 2 of Histone H2B type 2-F | 23 | 3 | 16 |
| HIST1H4A | P62805 | Histone H4 | 42 | 5 | 19 |
| HIST1H3A | P68431 | Histone H3.1 | 10 | 2 | 4 |
| HIST1H1D | P16402 | Histone H1.3 | 11 | 5 | 9 |

**Supplementary Table 1: Histone proteins identified in urine using mass spectrometry.**

| **Sample Number** | **Cancer Type** | **Primary Cancer Site** | **Cancer Type** | **Cancer Stage** | **Sample ID** |
|---|---|---|---|---|---|
| 31 | Adult | Pancreas | Pancreatic Adenocarcinoma | I | Pancreatic_urine_4 |
| 32 | Adult | Pancreas | Pancreatic Adenocarcinoma | I | Pancreatic_urine_9 |
| 33 | Adult | Pancreas | Pancreatic Adenocarcinoma | II | Pancreatic_urine_10 |
| 34 | Adult | Pancreas | Pancreatic Adenocarcinoma | II | Pancreatic_urine_12 |
| 35 | Adult | Pancreas | Pancreatic Adenocarcinoma | II | Pancreatic_urine_2 |
| 36 | Adult | Pancreas | Pancreatic Adenocarcinoma | II | Pancreatic_urine_5 |
| 37 | Adult | Pancreas | Pancreatic Adenocarcinoma | II | Pancreatic_urine_8 |
| 38 | Adult | Pancreas | Pancreatic Adenocarcinoma | IV | Pancreatic_urine_1 |
| 39 | Adult | Pancreas | Pancreatic Adenocarcinoma | IV | Pancreatic_urine_11 |
| 40 | Adult | Pancreas | Pancreatic Adenocarcinoma | IV | Pancreatic_urine_3 |
| 41 | Adult | Pancreas | Pancreatic Adenocarcinoma | IV | Pancreatic_urine_6 |
| 42 | Adult | Pancreas | Pancreatic Adenocarcinoma | IV | Pancreatic_urine_7 |
| 43 | Pediatric Solid Tumor | Rib | Ewing Sarcoma | Localized | Pediatric_urine_9 |
| 44 | Pediatric Solid Tumor | Maxilla | Ewing Sarcoma | Localized | Pediatric_urine_1 |
| 45 | Pediatric Solid Tumor | Psoas | Ewing Sarcoma | Localized | Pediatric_urine_3 |
| 46 | Pediatric Solid Tumor | Rib | Osteosarcoma | Localized | Pediatric_urine_4 |
| 47 | Pediatric Solid Tumor | Femur | Osteosarcoma | Localized | Pediatric_urine_6 |
| 48 | Pediatric Solid Tumor | Femur | Osteosarcoma | Localized | Pediatric_urine_5 |
| 49 | Pediatric Solid Tumor | Kidney | Rhabdoid Tumor | III | Pediatric_urine_10 |
| 50 | Pediatric Solid Tumor | Neck | Rhabdomyosarcoma | Localized | Pediatric_urine_2 |
| 51 | Pediatric Solid Tumor | Kidney | Wilms Tumor | III | Pediatric_urine_7 |
| 52 | Pediatric Solid Tumor | Kidney | Wilms Tumor | II | Pediatric_urine_8 |

**Supplementary Table 2: Clinical characteristics of cancer patients.**
